# *NO GAMETOPHORES 2* is a novel regulator of the 2D to 3D growth transition in the moss *Physcomitrium patens*

**DOI:** 10.1101/2020.07.21.213728

**Authors:** Laura A. Moody, Steven Kelly, Roxaana Clayton, Zoe Weeks, David M. Emms, Jane A. Langdale

## Abstract

The colonization of land by plants was one of the most transformative events in the history of life on Earth. The transition from water, which coincided with and was likely facilitated by the evolution of 3-dimensional (3D) growth, enabled the generation of morphological diversity on land. In many plants, the transition from 2-dimensional (2D) to 3D growth occurs during embryo development. However, in the early divergent moss *Physcomitrium patens* (formerly *Physcomitrella patens*), 3D growth is preceded by an extended filamentous phase that can be maintained indefinitely. Here, we describe the identification of the cytokinin-responsive *NO GAMETOPHORES 2* (*PpNOG2*) gene, which encodes a shikimate o-hydroxycinnamoyltransferase. In mutants lacking *PpNOG2* function, transcript levels of *CLAVATA* and *SCARECROW* genes are significantly reduced, excessive gametophore initial cells are produced, and buds undergo premature developmental arrest. Our results suggest that PpNOG2 functions in the ascorbic acid pathway leading to cuticle formation, and that NOG2-related genes were co-opted into the lignin biosynthesis pathway after the divergence of bryophytes and vascular plants. We present a revised model of 3D growth in which PpNOG2 comprises part of a feedback mechanism that is required for the modulation of gametophore initial cell frequency. We also propose that the 2D to 3D growth transition in *P. patens* is underpinned by complex auxin-cytokinin crosstalk that is regulated, at least in part, by changes in flavonoid metabolism.

## INTRODUCTION

One of the key developmental innovations that facilitated the colonization of land by plants was the evolution of 3-dimensional (3D) growth. In water dwelling Charophyte algae, the closest living relatives of land plants, cells only cleave in one or two planes to produce filaments, mats or branches. By contrast, land plants produce apical initial cells that can cleave in three (or more) planes to produce the elaborate morphologies that shape the terrestrial landscape [1,2]. In the vast majority of land plants, the transition from 2- dimensional (2D) to 3D growth occurs during the development of the embryo. However, in the early divergent moss *Physcomitrium patens* (formerly known as *Physcomitrella patens*) [3], the 2D to 3D growth transition occurs twice during the life cycle; initially during the transition from 2D filamentous growth to the development of 3D gametophores (‘leafy shoots’), and then during the development of the embryo [4, 5]. *P. patens* represents an ideal model organism in which to genetically dissect 3D growth because disruption of the 2D to 3D growth transition does not cause lethality.

The *P. patens* life cycle begins with the germination of haploid spores, which produce apical initial cells that grow in 1-dimension (1D) to produce filaments of chloronemal cells and then caulonemal cells, which are collectively known as the protonema [6, 7]. Growth can also occur in 2D or 3D when side branch initials are produced that develop into either secondary 2D protonema (approx. 95%) or 3D gametophores (approx. 5%) [4, 5, 8]. Side branch initials that give rise to 3D gametophores are known as gametophore initial cells (or bud initials). Auxin and cytokinin are required both for the formation of gametophore initial cells and for the coordination of cell division processes within the developing buds [9-14]. A gametophore initial cell first divides obliquely to establish an apical and a basal cell, which then both divide obliquely and perpendicular to the first division plane. An additional two rotating divisions establish a tetrahedral cell, which can both self-renew and divide in three planes to produce a mature 3D gametophore bearing leaf-like phyllids arranged in a spiral phyllotaxy, along with both antheridia and archegonia. Gametes combine during fertilization, leading to the formation of a diploid sporophyte, which releases haploid spores to restart the life cycle [5].

Previous studies have highlighted a number of genes that are essential for the 2D to 3D growth transition in *P. patens*. For example, orthologues of the Arabidopsis genes *AINTEGUMENTA, PLETHORA* and *BABY BOOM* (*PpAPB1-4*) are necessary and sufficient for gametophore initial cell formation [8] whereas *DEFECTIVE KERNEL 1* (*PpDEK1*) [15-18] and *CLAVATA* [19] negatively regulate gametophore initial cell formation and positively regulate division plane orientation. In contrast, *NO GAMETOPHORES 1* (*PpNOG1*) acts as a positive regulator of both gametophore initial cell formation and division plane orientation in developing buds [20]. A current model for *P. patens* 3D growth regulation suggests that *PpDEK1* and *PpNOG1* act antagonistically to regulate the transcription of the *PpAPB* genes, which switch on a transcriptional network that facilitates gametophore initial cell formation. Correct division plane orientation in the developing gametophore is then regulated through a feedback circuit that comprises *PpNOG1, PpDEK1* and *CLAVATA* [20, 21]. Here, we describe the identification of a further regulator of 3D growth in *P. patens*. The discovery of *NO GAMETOPHORES 2* (*PpNOG2*), which encodes a shikimate o-hydroxycinnamoyltransferase, enables the role of auxin-cytokinin crosstalk to be incorporated into the regulatory pathway.

## RESULTS

### *Ppnog2-R* mutants produce supernumerary buds but fail to specify 3D growth

A forward genetic screen of 9,000 UV-mutagenized lines of the Villersexel (Vx::mCherry) [22] strain of *P. patens*, identified two mutants that exhibit normal 2D filamentous tip growth and branching but fail to transition to 3D growth. In contrast to *P. patens no gametophores 1 – Reference* (*Ppnog1-R*), where gametophore initials are suppressed [20], excessive numbers of gametophore initial cells (up to eight times the frequency of wild type) are produced in *no gametophores 2 – Reference* (*Ppnog2-R)* mutants (**Figure 1**). Despite the formation of supernumerary initial cells, no gametophores are produced in *Ppnog2-R* mutants because aberrant cell division patterns prevent the formation of a tetrahedral apical cell. In wild type *P. patens*, the first division of the gametophore initial cell is roughly perpendicular to the parent caulonemal cell from which the initial is derived, producing the apical and basal cells (**Figure 2A**). The apical and basal cells then divide obliquely and perpendicular to the first division plane (**Figures 2B and 2C**), with subsequent divisions enabling the establishment of the apical cell (**Figure 2D**). In *Ppnog2-R* mutants, the first oblique division of the gametophore initial cell is correctly oriented (**Figure 2E**), as is the plane of the second division (**Figure 2F**). However, a range of developmental abnormalities occur thereafter. In some cases, the apical initial cell maintained stemness, but then failed to divide in the correct manner (**Figure 2G-I**). In other cases, the newly formed cell acquired an apical cell identity and expanded excessively to generate a bud that was bilaterally symmetrical (**Figure 2J, K**). The pre-defined apical initial cell and the newly expanded apical cell then entered a concurrent cell division programme to produce ‘bifurcating’ twin ‘buds’ that arrested early during development (**Figure 2L**). Without exception, the aberrant cell division patterns exhibited by *Ppnog2-R* mutants prevented the formation of gametophores. The two distinct phenotypes in *Ppnog2-R* mutants, (formation of supernumerary initials and impaired cell division plane orientation) imply that *PpNOG2* negatively regulates gametophore initial cell formation and positively regulates the establishment of 3D growth through correct specification of the gametophore apical cell.

**Figure 1.**
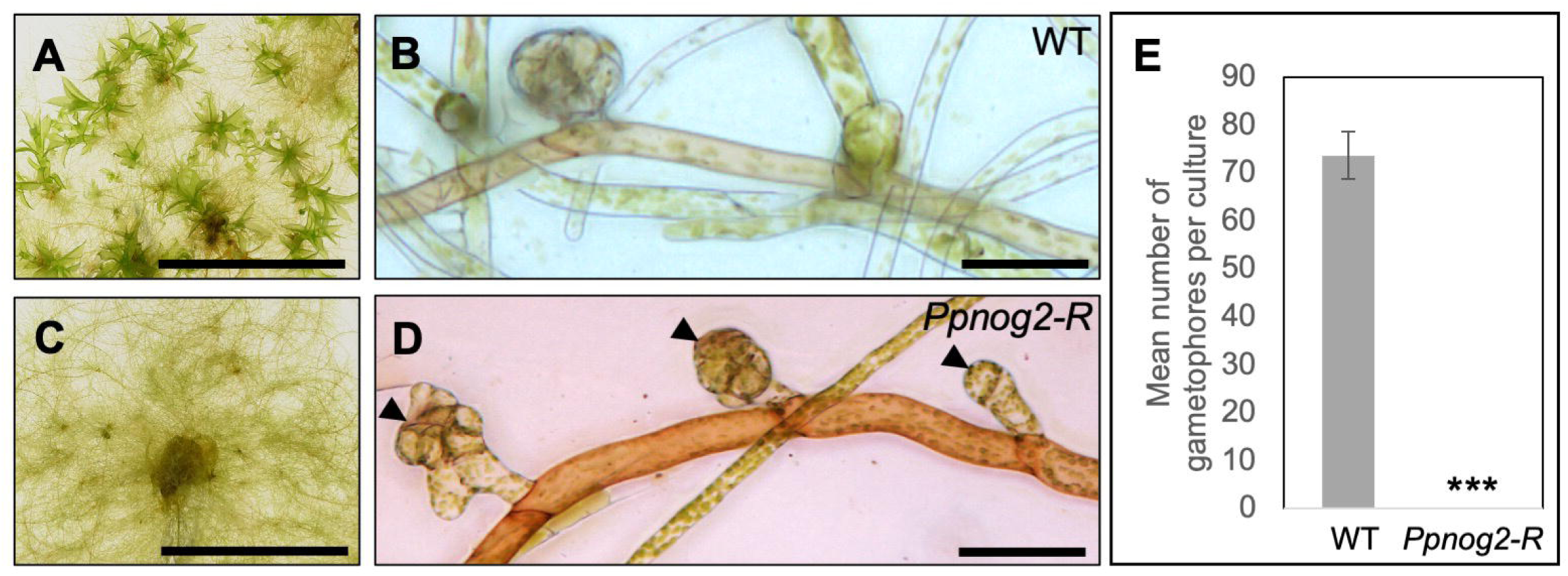
*Ppnog2-R* mutants produce supernumerary buds. (A-D) 1 month old Vx::mCherry (WT) (A and B) and *Ppnog2-R* (C and D) plants showing the presence (WT, A) and absence (*Ppnog2-R*, C) of gametophores and the protonemal filaments (B and D) with supernumerary buds visible in *Ppnog2-R* (D). (E) Mean number of gametophores per culture (n = 10) ± SEM (WT = 73.7 ± 4.90; Ppnog2-R = 0 ± 0; t test p < 0.05 ***). Scale bars, 1 cm (A and C) and 50 µm (B and D).

**Figure 2.**
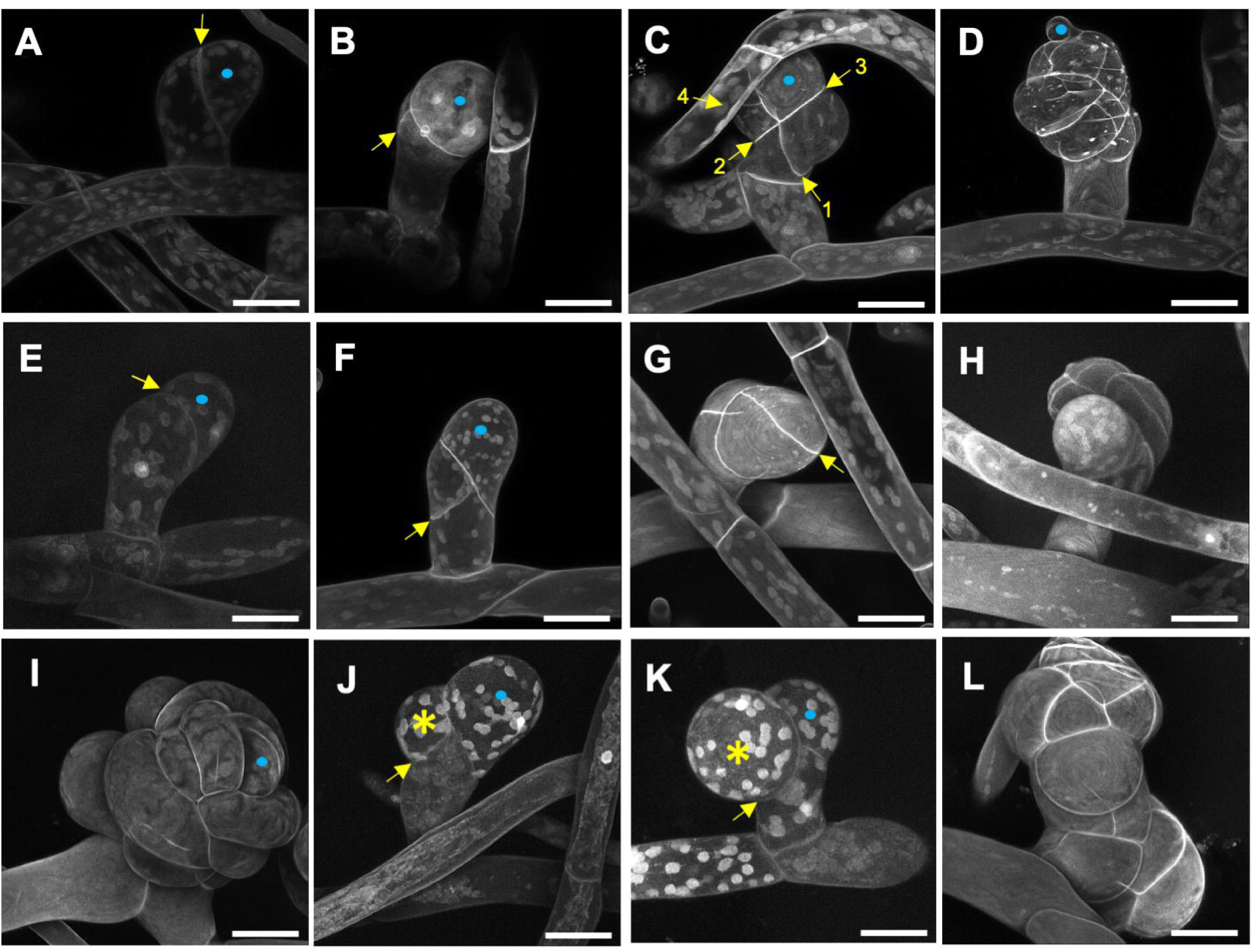
*Ppnog2-R* mutants fail to specify 3D growth. (A-L) Propidium Iodide stained Vx::mCherry (WT) (A-D) and *Ppnog2-R* (E-L) buds at 2 (A and E), 3 (B, F, J and K), 4/5 (C and G) and 5+ (D, H, I and L) cell stages. Scale bars = 25 µm; • = gametophore initial. Yellow arrows indicate the most recent division in each bud. Yellow asterisks (*) denote cell in *Ppnog2-R* (J and K) that undergoes abnormal swelling.

### The *PpNOG2* gene encodes a SHIKIMATE O-HYDROXYCINNAMOYLTRANSFERASE

To identify the causative mutation in *Ppnog2-R* mutants, segregating populations were derived from diploid somatic hybrids generated between the infertile *Ppnog2-R* mutant and fertile lines of the Gransden (Gd::GFP) strain [22] (**Figure 3A-C, Figure S1A**). The observed phenotypic segregation ratio of progeny was consistent with meiosis from a tetraploid (**Figure 3D**), providing mutant and wild-type individuals for gene identification by genome sequencing. Genomic DNA was prepared from 120 mutant individuals (no 3D growth) and pooled in equimolar amounts to form the ‘mutant pool’ and similarly from 120 wild type individuals (normal 3D growth) to form the ‘wild type pool’. Mutant and wild type pools were then sequenced at 30X coverage. The parental Vx::mCherry and Gd::GFP lines were sequenced previously [23]. To identify the genetic interval containing the *Ppnog2-R* mutation, mutant allele frequencies were plotted across all 27 chromosomes in the *P. patens* genome assembly for 120 mutant individuals and 120 wild type individuals. This revealed a mutant allele frequency of 1 on chromosome 2 in the mutant pool (**Figure S1B**), and a corresponding mutant allele frequency of 0.4 in the wild-type pool (**Figure S1C**). A comprehensive analysis of single nucleotide polymorphisms (SNPs), insertions and deletions within the defined genetic interval revealed mutations affecting five different annotated genes (**Figure 3E**). The most likely cause of the *Ppnog2-R* mutant phenotype was a mutation in either Pp3c2_27620 or Pp3c2_29140 because the transcripts of these mutated genes contained premature termination codons. The mutation in Pp3c2_27620 results in a nonsynonymous nucleotide substitution (G^82^G), but the frameshift mutation leads to the introduction of a stop codon in the downstream sequence. Pp3c2_27620 encodes a short protein composed of 106 amino acid residues, with no clearly identifiable domain sequences. A G>A transition generated a premature stop codon (W^541^Ter) in Pp3c2_29140, which encodes a shikimate O-hydroxycinnamoyltransferase containing two transferase domains.

**Figure 3.**
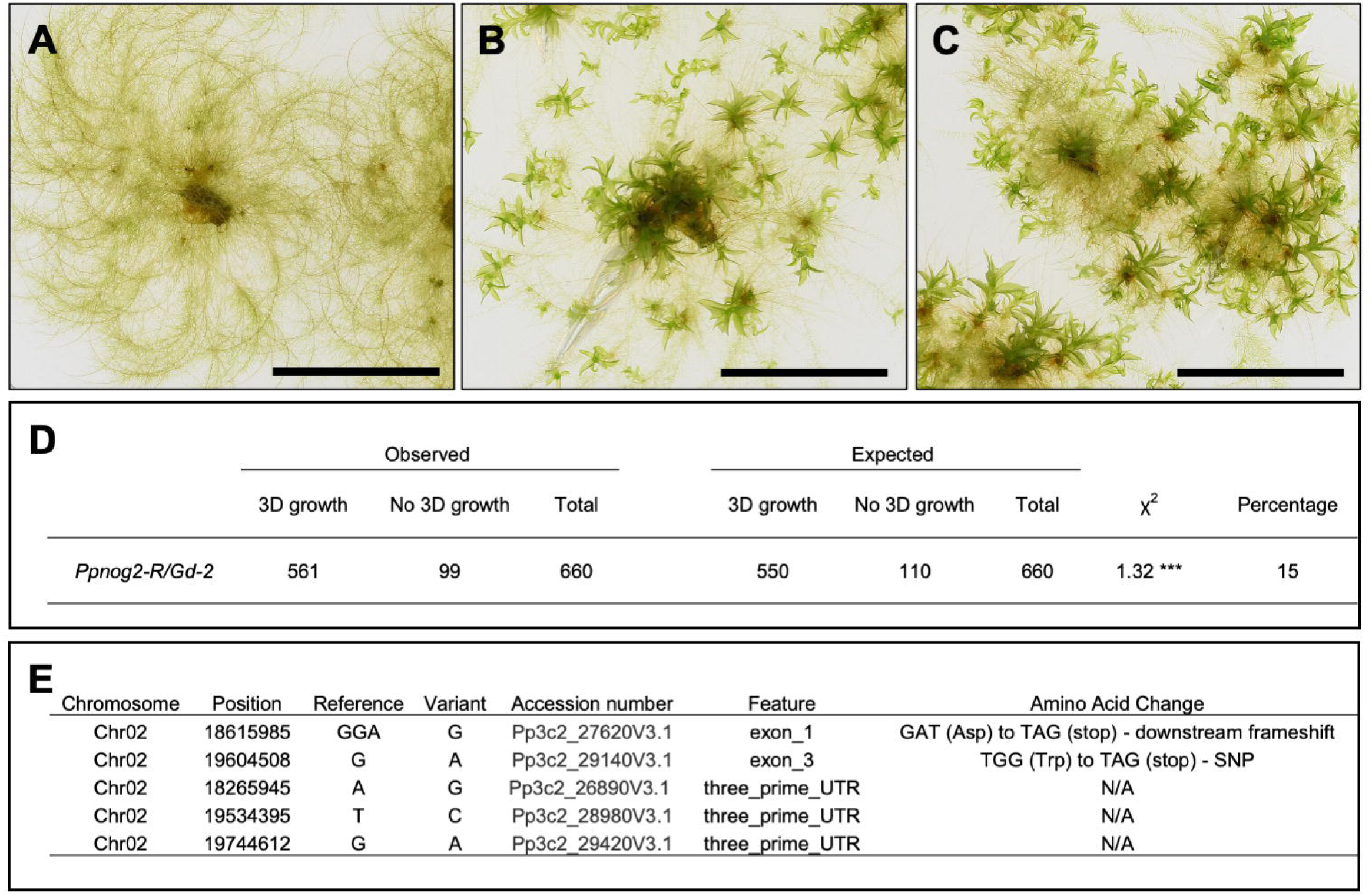
Identification of the *PpNOG2* locus by somatic hybridization combined with bulk segregant analysis. (A-C) Representative images of 1-month old Ppnog2-R plant without gametophores (A), Vx::mCherry plant with gametophores (B) and Ppnog2-R/Gd::GFP hybrid with gametophores (C). Scale bars = 1 cm. (D) Phenotypic analysis of spore progeny derived from a single Ppnog2-R/Gd::GFP hybrid line (no. 2). Numbers are consistent with meiosis from a tetraploid sporophyte. Chi-square P<0.05 ***. E) Candidate genes on Chromosome 2 within the genetic interval containing the *PpNOG2* genetic locus. See also **Figure S1**.

To determine whether mutations in Pp3c2_27620 or Pp3c2_29140 cause the *Ppnog2-R* mutant phenotype, we first tried to amplify both transcripts from wild type and *Ppnog2-R* cDNA. Attempts to amplify Pp3c2_27620 from cDNA were not successful, but the Pp3c2_29140 transcript was isolated with ease. The wild type version of the Pp3c2_29140 transcript was identical to that derived from the v3.3 annotation of the v3.0 *P. patens* genome assembly. However, two different splice variants of the Pp3c2_29140 transcript were detected in the *Ppnog2-R* mutant (**Figure 4A**). The alternatively spliced transcripts lead to one (v1) or both (v2) of the transferase domains being deleted from the encoded protein (**Figure 4B**) suggesting that the *Ppnog2-R* mutant phenotype is not caused by the premature stop codon as such, but by nonsense-associated alternative splicing limiting the translation of a potentially harmful truncated protein [24]. Intriguingly the same phenomenon was observed in the *Ppnog1-R* mutant [20]. Collectively, these data suggested that a mutation in the Pp3c2_29140 gene was responsible for the *Ppnog2-R* phenotype. To validate this hypothesis, the rice actin promoter was used to drive the expression of the full-length wild-type coding sequence in the *Ppnog2-R* mutant (**Figure S2**). The mutant phenotype was fully complemented by the Pp3c2_29140 gene (**Figure 4C, D**), and therefore the gene was named *NO GAMETOPHORES 2* (*PpNOG2*).

**Figure 4.**
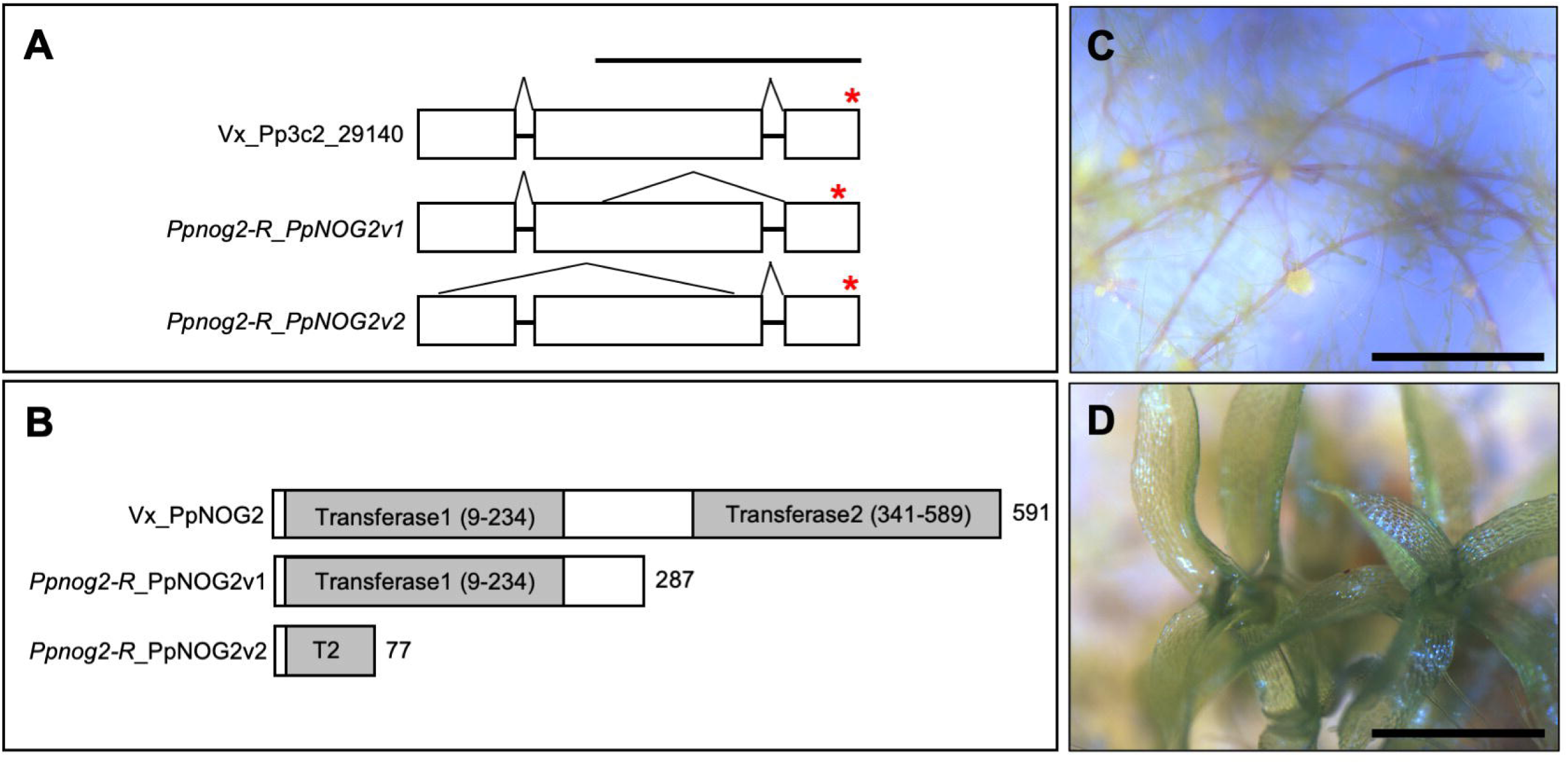
The *PpNOG2* gene encodes a SHIKIMATE O- HYDROXYCINNAMOYLTRANSFERASE. *(A) PpNOG2* transcripts in Vx::mCherry (Vx_Pp3c2_29140) and *Ppnog2-R* (v1 and v2). Exons (blocks) are separated by introns (lines); splicing patterns and in-frame stop codons (*) are indicated. Scale = 1kb. (B) PpNOG2 protein in Vx::mCherry (Vx_PpNOG2) and both variants of PpNOG2 protein in the *Ppnog2-R* mutant. *Ppnog2-R*_PpNOG2v1 contains an intact Transferase 1 (T1) domain but Transferase 2 (T2) is absent. In *Ppnog2- R*_PpNOG2v2, T1 is absent and T2 is heavily truncated. The number of amino acid residues is indicated to the right of each image. (C, D) Representative images of 1-month old *Ppnog2-R* (C) and *Ppnog2-R* complemented with pAct::*PpNOG2*-mGFP (D). Scale bars = 1 mm. See also **Figure S2-S4** and **Data S2**.

To investigate whether PpNOG2 contains transmembrane (TM) domains, typical of other transferase proteins in plants, the PpNOG2 protein sequence was scanned for the presence of putative membrane-spanning regions using TMpred [25]. Three TM domains were predicted to exist in the PpNOG2 protein. The most well supported model indicated that the N-terminal end of PpNOG2 resides within the cytosol, and that the C-terminal end is located on the external surface of the membrane. The model also indicated the presence of a significant extracellular loop region. Crucially, the highly conserved aspartic acid and histidine residues that are putative proton acceptors (‘active sites’) required for transferase activity [26] are present in PpNOG2 (**Figure S3, Data S1**). To validate the localization of PpNOG2 *in planta*, the rice actin promoter was used to drive the expression of a PpNOG2- GFP fusion protein in *P. patens* protoplasts. Consistent with *in silico* predictions, confocal microscopy revealed that PpNOG2-GFP localizes at or near to the plasma membrane (**Figure S3**).

### PpNOG2 orthologues function in the lignin biosynthesis pathway

To infer phylogenetic relationships for PpNOG2, orthologous sequences were retrieved from the genomes of *P. patens*, the liverwort *Marchantia polymorpha*, the lycophyte *Selaginella moellendorfii* and the flowering plants *Oryza sativa* and *Arabidopsis thaliana*. Aligned amino acid sequences were used to infer a consensus tree, which was rooted on related sequences from chlorophyte green algae. The topology of the tree revealed that the Arabidopsis orthologue of *PpNOG2* is hydroxycinnamoyl-CoA:shikimate hydroxycinnamoyl transferase (HCT; At5g48930) (**Figure S4**). In angiosperms, HCT first catalyses the transfer of alcohols or amines to p-coumaroyl CoA to produce p-coumaroyl shikimate, which subsequently acts as a substrate for CYP98. CYP98 converts p-coumaroyl shikimate into caffeoyl shikimate, which then acts as an additional substrate for HCT. Subsequent downstream steps of the pathway lead to lignin biosynthesis [27]. PpNOG2 orthologs thus play important roles in phenylpropanoid biosynthesis and in the formation of lignin in vascular plants.

The structural features and localization pattern of PpNOG2 are consistent with a functional HCT protein but it has been reported that monolignols are not present in bryophytes, and that the HCT substrate caffeoyl shikimate is undetectable in *P. patens* [28]. It has been shown, however, that the transfer of alcohols or amines to p-coumaroyl CoA by *P. patens* HCT (PpNOG2) produces p-coumaroyl threonate rather than the p-coumaroyl shikimate produced by HCT in Arabidopsis. Interestingly, both the formation of p-coumaroyl threonate by PpNOG2, and subsequent modification by PpCYP98A3, occur in an ascorbic acid pathway, suggesting that these enzymes were co-opted into the lignin biosynthesis pathway after the divergence of bryophytes and vascular plants.

In angiosperms, p-Coumaroyl-CoA lies at the intersection between flavonoid and lignin biosynthesis. Consequently, disruption of HCT function can divert the metabolic flux into an overaccumulation of flavonoids, which can inhibit auxin transport [29]. In *P. patens*, the inhibition of auxin transport by naphthylphthalamic acid (NPA) increases sensitivity of wild- type plants to auxin at low concentrations and disrupts 3D growth. Although apical cell identity is maintained, buds undergo premature developmental arrest and resemble those treated with auxin at high concentrations. The authors suggest that a functional auxin transport system is necessary to divert exogenously applied auxin and prevent unfavourable accumulation in cells [30]. Interestingly, developing buds of the *Ppnog2-R* mutant share remarkable similarities with those affected in auxin homeostasis (**Figure 2I**).

The initial step of flavonoid biosynthesis is catalysed by chalcone synthases, which are known to be functional in *P. patens* [31-33]. As such we hypothesize that blocking metabolic flux into the ascorbic acid pathway in which both *PpNOG2* and *PpCYP98A3* play a role, would lead to a similar overaccumulation of flavonoids and perturbation of auxin homeostasis. This hypothesis is consistent with the *Ppnog2-R* mutant phenotype, with the observation that loss of PpCYP98A3 function leads to the formation of gametophores that undergo early developmental arrest [28, 34], and the known role of auxin in gametophore development [30, 35].

### *PpNOG2* acts downstream of *PpNOG1* but upstream of *CLAVATA*

We previously proposed a model for 3D growth in *P. patens*, in which PpNOG1 and PpDEK1 operate in an antagonistic manner to regulate the expression of *PpAPB* genes. Auxin and cytokinin response regulators, the transcriptional targets of *PpAPB* genes, then enable the formation of gametophore initial cells. Subsequently, PpNOG1, PpDEK1 and CLAVATA act within the gametophore initial to ensure that cell division planes are correctly oriented to produce a tetrahedral initial cell that can facilitate 3D growth (**Figure S5**) [20, 21]. To understand where *PpNOG2* fits into this pathway, quantitative RT-PCR experiments were carried out to determine transcript accumulation profiles of known 3D regulators in wild type and in *Ppnog1-R* and *Ppnog2-R* mutants.

To place *PpNOG2* function in the context of *PpNOG1*, which positively regulates both gametophore initial cell formation and division plane positioning [20], reciprocal quantitative RT-PCR experiments were performed. *PpNOG1* transcript accumulation was unaffected in the *Ppnog2-R* mutant (**Figure 5A**) but *PpNOG2* transcript accumulation was dramatically reduced in the *Ppnog1-R* mutant compared to wild type (**Figure 5B)**. *PpNOG1* function therefore induces the accumulation of *PpNOG2* transcripts (either directly or indirectly).

**Figure 5.**
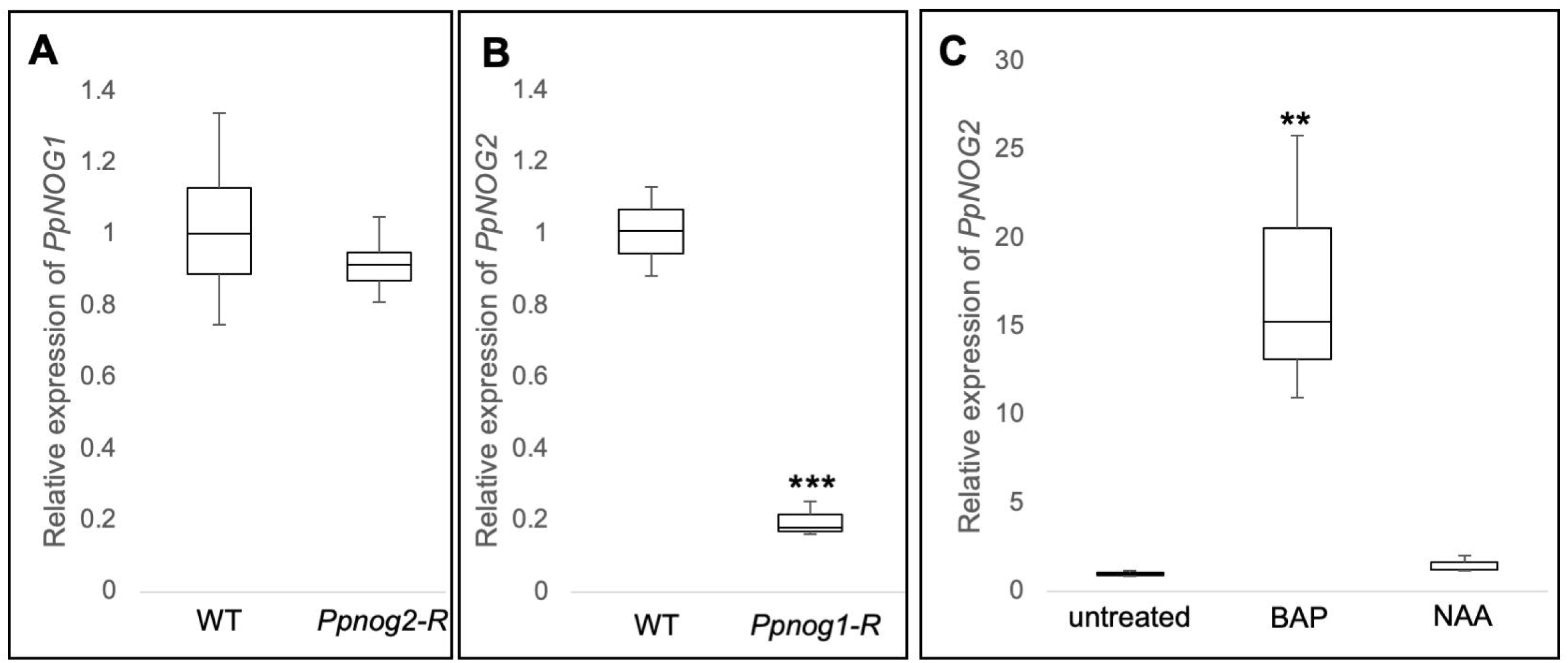
*PpNOG2* acts downstream of *PpNOG1* and is regulated by cytokinin. (A-C) Relative transcript levels in 12 d old protonemata: *PpNOG1* in WT and *Ppnog2-R* (A), *PpNOG2* in WT and *Ppnog1-R* (B), *PpNOG2* in wild-type ± BAP or NAA (C). ANOVA, **, p<0.01; ***, p<0.001. See also **Figure S5**.

The supernumerary buds formed in the *Ppnog2-R* mutant resemble those of wild type protonema treated with exogenous cytokinin, but also resemble those affected in auxin homeostasis [30]. We hypothesise that *PpNOG2* negatively regulates bud initiation by promoting auxin signalling. To investigate whether *PpNOG2* transcript abundance is regulated by cytokinin or auxin, transcript levels were quantified in 1-week old wild type protonemata that had been grown in the presence or absence of the cytokinin analogue 6- Benzylaminopurine (BAP) or the auxin analogue 1-Naphthaleneacetic acid (NAA). Transcript levels of *PpNOG2* were dramatically induced by exogenous BAP treatment whereas they were completely unaffected by NAA treatment (**Figure 5C**). The accumulation of *PpNOG2* transcripts is thus induced by cytokinin.

Transcript levels of the four *PpAPB* transcription factors that are necessary and sufficient to drive gametophore formation [8] are significantly reduced in *Ppnog1* disruption mutants that produce fewer gametophore initial cells than wild type [20]. To investigate whether *PpAPB* transcript levels are correspondingly increased in *Ppnog2-R* mutants that produce excessive gametophore initial cells, qRT-PCR was carried out using cDNA isolated from 12 d old wild type and *Ppnog2-R* mutant protonemata. Only *PpAPB1* transcript levels were significantly increased in *Ppnog2-R* mutants compared to wild type. *PpAPB2* and *PpAPB4* transcript levels were also increased in the mutant and *PpAPB3* transcript levels decreased, but none of these differences were significant (**Figure 6A**). Neither the pattern nor level of *PpAPB* gene expression correlated with the number of gametophore initial cells formed in the *Ppnog2-R* mutant, and as such any differences from wild-type most likely represent indirect effects of altered stem cell fate. *PpNOG2* thus operates downstream of the *PpAPB* genes.

**Figure 6.**
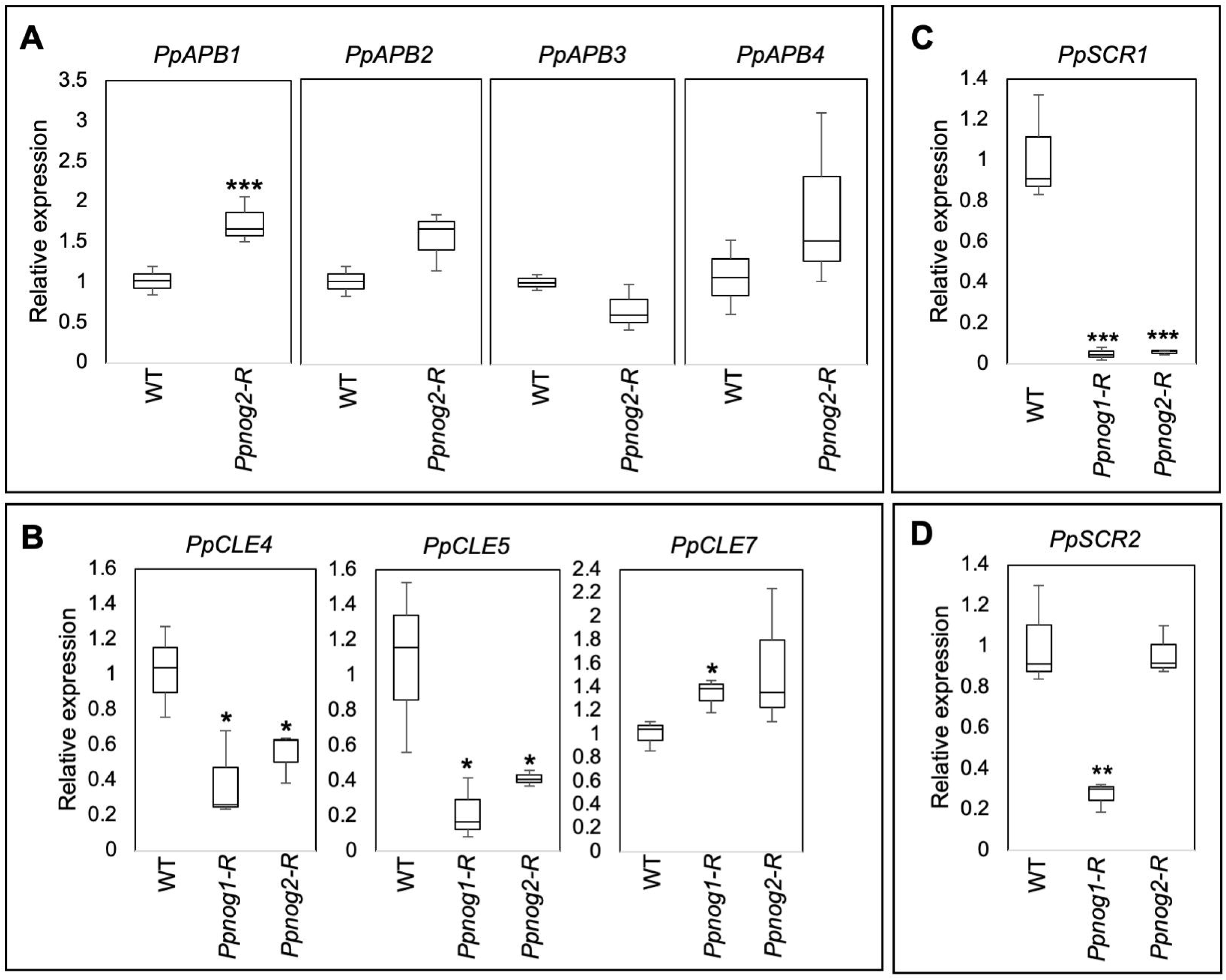
PpNOG2 act upstream of *CLAVATA* and *SCARECROW* genes. (A-D) Relative transcript levels in 12 d old protonemata: *PpAPB1-4* in Vx::mCherry (WT) and *Ppnog2-R* (A); *PpCLE4, PpCLE5* and *PpCLE7* in WT, and in *Ppnog1-R* and *Ppnog2-R* mutants (B); *PpSCR1* (C) and *PpSCR2* (D) in WT, and in *Ppnog1-R* and *Ppnog2-R* mutants. Statistical analysis: t-test, P<0.05 *** (A); ANOVA, p<0.01 **; p<0.001 *** (B-D). See also **Figure S6**.

The overbudding phenotype exhibited by *Ppnog2-R* mutants bears a closer resemblance to those of *Ppdek1, Ppcle, Ppclv* and *Pprpk2* disruptant mutants, than to the *Ppnog1-R* mutant [15-19]. Phenotypic observations alone suggest that *PpNOG1* and *PpDEK1* act at the first step of the pathway that initiates the 2D to 3D growth transition. Using the same logic, it would be reasonable to suggest that *PpNOG2*, and then *CLAVATA* operate sequentially at subsequent stages of development. To better understand the relationships between these different players, transcript accumulation profiles of *PpDEK1* and *CLAVATA* (*PpCLE1-7, PpCLV1A, PpCLV1B* and *PpRPK2*) genes were compared in 12 d old wild type protonemata and in both *Ppnog1-R* and *Ppnog2-R* mutants. Of the 11 genes examined (see methods), only three showed transcript levels that were significantly different between wild-type and either *Ppnog1-R* or *Ppnog2-R* mutants. *PpCLE4* and *PpCLE5* levels were significantly reduced in both *Ppnog1-R* and *Ppnog2-R* mutants compared to wild type, whereas *PpCLE7* levels were significantly increased in *Ppnog1-R* but not in *Ppnog2-R* mutants (**Figure 6B**). *PpCLE7* transcript levels are thus suppressed by PpNOG1, whereas PpCLE*4* and *PpCLE5* levels are induced by both *PpNOG1* and *PpNOG2*.

### SCARECROW transcription factors are markers of 3D growth

The first visual marker of the 3D growth transition in *P. patens* is an asymmetric cell division in the gametophore initial cell. To identify molecular markers of this event, we took a candidate gene approach, selecting genes based on their documented gene expression profiles in *P. patens* [36] and their known roles in asymmetric cell divisions in angiosperms. Transcript accumulation profiles were quantified in 12 d old protonemata of wild-type, plus both *Ppnog1-R* and *Ppnog2-R* mutants, for *SCARECROW 1* (*PpSCR1*), *SCARECROW 2* (*PpSCR2*) and *SCARECROW 3* (*PpSCR3*). The *PpSCR3* transcript were not detected in 12 d old protonemata and were therefore not investigated further. However, quantification of *PpSCR1* transcript levels revealed a significant reduction in both *Ppnog1-R* and *Ppnog2-R* mutants (**Figure 6C**) whereas levels of *PpSCR2* transcript were significantly reduced in the *Ppnog1-R* mutant but not in the *Ppnog2-R* mutant (**Figure 6D, Figure S6**).

It has been reported that SCR genes regulate auxin-cytokinin crosstalk in angiosperms [37, 38]. Given the opposing bud initiation phenotypes in *Ppnog1-R* and *Ppnog2-R* mutants, our results suggest that *PpSCR1* and *PpSCR2* modulate auxin-cytokinin crosstalk at different stages of the 2D to 3D growth transition, both downstream of *PpNOG1* (*PpSCR2*) and downstream of *PpNOG2* (*PpSCR1*).

### A revised model for 3D growth regulation in *P. patens*

Results reported here add a new dimension to current models for the 3D growth transition in *P. patens* (**Figure 7**). As reported previously, we propose that PpNOG1 acts antagonistically to PpDEK1 at the earliest stage of 3D growth initiation to induce the degradation of a repressor of *PpAPB* activity. The subsequent activation of cytokinin biosynthetic genes, proposed targets of the *PpAPB* genes, enables the formation of gametophore initial cells. Based on the findings reported here, we propose that PpNOG2 comprises part of a feedback mechanism that is required for the modulation of gametophore initial cell frequency. Our results suggest that the 2D to 3D growth transition is underpinned by complex auxin-cytokinin crosstalk that may be regulated, at least in part, by changes in flavonoid metabolism. In response to elevated cytokinin levels, we propose that PpNOG2 activates an auxin response that is dependent on CLAVATA signalling. The correct modulation of local auxin and cytokinin concentrations ensure that division planes within developing buds are correctly oriented, and that unnecessary bud initiation is eliminated.

**Figure 7.**
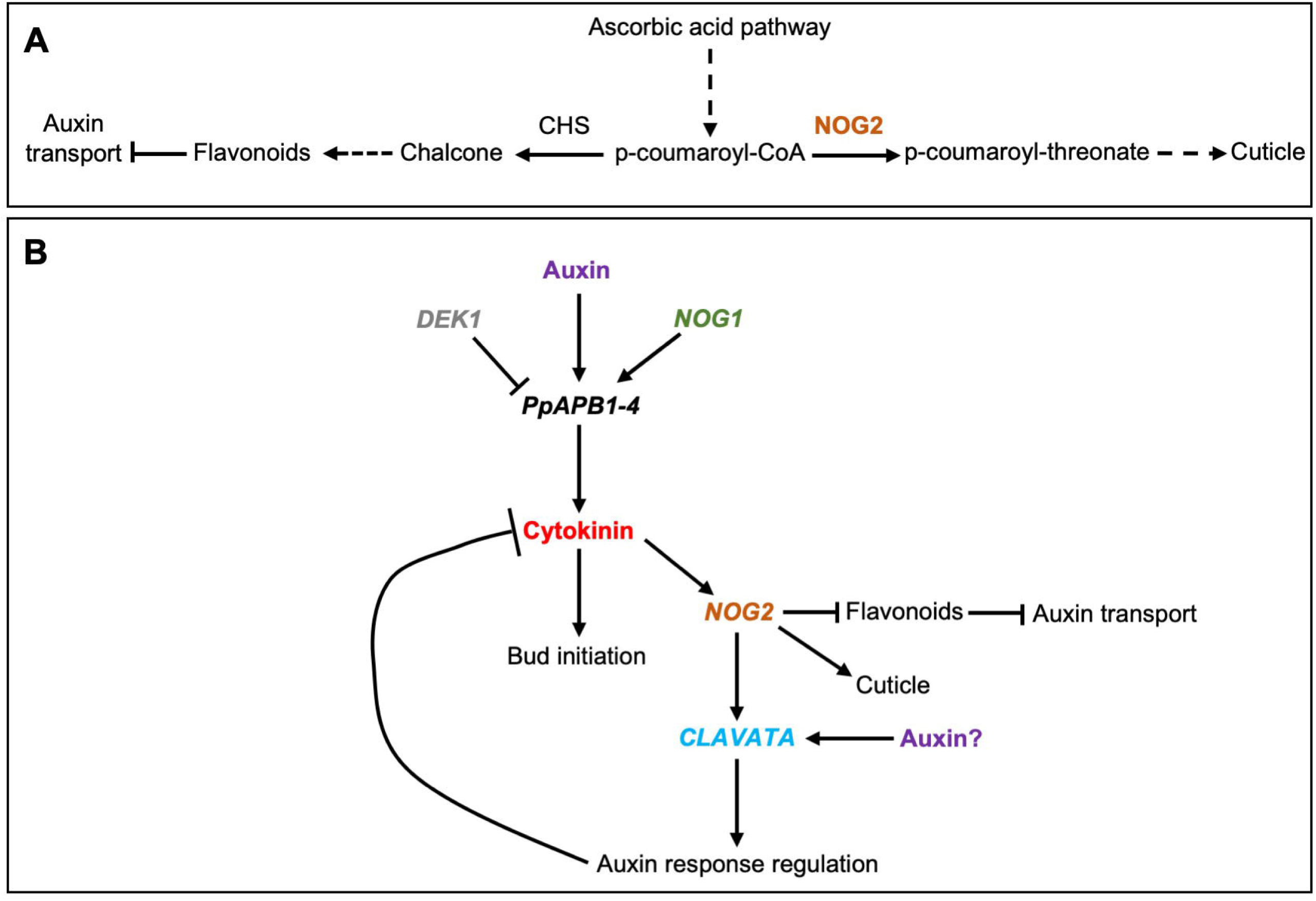
Model for 3D growth regulation in *P. patens*. A) Proposed involvement of PpNOG2 in the ascorbic acid pathway leading to cuticle formation. When PpNOG2 is absent, metabolic flux is directed through to flavonoid biosynthesis through enhanced chalcone synthase (CHS) activity. Increased flavonoids cause inhibition of auxin transport and suppression of the auxin response. B) Model for PpNOG2 function in the context of 3D growth regulation: PpNOG1 and PpDEK1 operate antagonistically to induce the degradation of a repressor of *PpAPB* activity. Activation of cytokinin biosynthesis initiates bud development and induces *PpNOG2* transcription. PpNOG2 then modulates auxin-cytokinin crosstalk, which is dependent on CLAVATA signalling. This ensures that further bud development is prevented, and that division planes within developing buds are correctly oriented.

## ACKNOWLEDGEMENTS

We are grateful to Pierre-François Perroud for providing the Vx::mCherry and the Gd::GFP marker lines; Yuji Hiwatashi for providing the pZAG1 plasmid; and Andrew Cuming for advice on UV-mediated mutagenesis. The work was funded by BBSRC (BB/M020517/1) grants (to J.A.L.) and by Royal Society University Research Fellowships to L.A.M. (URF\R1\191310) and S.K. (UF140484).

## AUTHOR CONTRIBUTIONS

L.A.M. conducted the experiments, with technical assistance from R.C; L.A.M., S.K., and J.A.L. conceived and designed the study; S.K. carried out the bioinformatics; Z.W., D.E., and J.A.L. assisted L.A.M. with Phylogenetic tree construction; and L.A.M. and J.A.L. wrote the manuscript.

## DECLARATION OF INTERESTS

The authors declare no competing interests.

## METHODS

### CONTACT FOR REAGENT AND RESOURCE SHARING

Further information and requests for resources and reagents should be directed to and will be fulfilled by the Lead Contact, Laura Moody. Please note that the transfer of transgenic *P. patens* lines will be governed by an MTA and will be dependent on appropriate import permits being acquired by the receiver.

### EXPERIMENTAL MODEL AND SUBJECT DETAILS

#### Plants

Vx::mCherry and Gd::GFP marker lines were obtained from Pierre-François Perroud [22], and the *Ppnog1-R* line was generated previously [20]. Plants were propagated under sterile conditions on BCD or BCDAT medium. BCD medium contained 250mg/L MgSO_4_.7H_2_O, 250mg/L KH_2_PO_4_ (pH6.5), 1010mg/L KNO_3_, 12.5mg/L FeSO_4_.7H_2_O, 0.001% Trace Element Solution (TES – 0.614mg/L H_3_BO_3_, 0.055mg/L AlK(SO_4_)_2_.12H_2_O, 0.055mg/L CuSO_4_.5H_2_O, 0.028mg/L KBr, 0.028mg/L LiCl, 0.389mg/L MnCl_2_.4H_2_O, 0.055mg/L CoCl_2_.6H_2_O, 0.055mg/L ZnSO_4_.7H_2_O, 0.028mg/L KI and 0.028mg/L SnCl_2_.2H_2_O) and 8g/L agar, supplemented with 1 mM CaCl_2_. BCDAT medium was additionally supplemented with 1 mM ammonium tartrate [39]. Plants were grown at 24 °C with a 16 h: 8 h, light (300mmol m^-2^ s^-1^): dark cycle. To induce sporophytes, tissues were grown for 3-6 months on peat plugs at 16°C with an 8 h: 16 h, light (300mmol m^-2^ s^-1^): dark cycle. Sporangia were sterilized in 20% Parozone bleach for 15 min at room temperature and washed three times in sterile water. Sporangia were ruptured using a sterile pipette tip, and the released spores were plated onto cellophane overlaid BCDAT medium and grown at 24 °C with a 16 h: 8 h, light (300mmol m^-2^ s^-1^): dark cycle.

## METHOD DETAILS

### UV mutagenesis and screening

1% Driselase was prepared in 8% mannitol and incubated for 15 min at room temperature. The solution was centrifuged for 3 min at 3,300 xg and then filter sterilized without disrupting the pellet. 7-day old Vx::mCherry protonemata was harvested from cellophane overlaid BCDAT medium, added to the 1% Driselase solution and incubated, with gentle agitation, for 40 min. The cell suspension was then filtered through a 40 µm cell strainer into a round bottomed tube and centrifuged for 3 min at 120 xg at room temperature with no braking. The pelleted cells were washed twice with 2 × 6 ml 8% mannitol, and then resuspended in 6 ml 8% mannitol. Cell density was determined using a haemocytometer and adjusted to a density of 5×10^4^ cells mL^-1^. 1 mL cell suspension was plated onto cellophane overlaid BCDATG medium (BCDAT supplemented with 10 mM CaCl_2_, 6% mannitol and 0.5% glucose). Plates containing protoplasts were administered a dose of UV radiation using a Stratalinker UV Crosslinker (75000 mJ) and then incubated at 24°C in the dark for 48 h to prevent photoactivatable DNA damage repair. Regenerating protoplasts were incubated in standard light conditions for one week. Cellophane discs were then transferred to BCDAT medium and grown for one week until regenerating protoplasts were clearly visible. Regenerating protoplasts were transferred to individual wells of 24-well plates containing BCD medium and incubated under standard growth conditions until ready for phenotypic analyses (typically two months).

### Microscopy

Fluorescence microscopy was carried out using a Leica SP5 confocal microscope. A 63x water immersive lens was used for all imaging. Propidium iodide (PI) was excited at 488 nm with 30% laser power and detected with at 600-630 nm. Tissues were submerged in 10 µgmL^-1^ PI for 1-5 min and then mounted on a slide in a drop of water. All other images were captured using a Leica DMRB microscope or a Leica M165C stereomicroscope, equipped with a QImaging Micropublishing 5.0 RTV camera.

### Somatic hybridization and segregation analysis

Protoplasts were isolated from both *Ppnog2-R* and Gd::GFP as described above, and adjusted to a cell density of 1×10^6^ cells mL^-1^ in 8% mannitol. 1×106 cells of each line were combined in a single round bottomed tube and mixed gently. Protoplasts were pelleted after centrifugation for 3 min at 120xg at room temperature with no braking, and then resuspended in 250 µl protoplast wash (PW) solution (10 mM CaCl_2_ and 8.5% mannitol) and 750 µl PEG/F (5 mM CaCl2 and 50% PEG 6000 (w/v)), and left to incubate at room temperature for 40 min. 1.5 mL of PW solution was then added to the tubes, mixed gently and left to incubate at room temperature for 10 min. An additional 10 mL PW solution was added to the cell suspension, and left to incubate for a further 10 min. This step was repeated. Protoplasts were then centrifuged at 120xg for 3 min at room temperature with no braking. The supernatant was removed, and the pellet resuspended in 3 mL 8% mannitol prior to plating 1 mL onto each of three cellophane overlaid petri dishes containing BCDATG medium. Plates were incubated in the dark for 48 h and then transferred to standard growth conditions. After one week, cellophane discs were removed and then transferred to BCDAT medium supplemented with both 50 µgmL^-1^ G418 and 20 µgmL^-1^ hygromycin B, to select for stable *Ppnog2-R*/Gd::GFP diploid hybrids (adapted from [40]).

Sporophyte induction was performed as described above. Spores were germinated on cellophane overlaid BCDAT medium for two weeks. Regenerating sporelings were then transferred to individual wells of 24-well tissue culture plates containing BCD medium, and grown for two months under standard growth conditions. Phenotypic characterisation of the lines was then carried out. 120 of the lines that failed to initiate 3D growth contributed to the mutant pool, and 120 of the lines could initiate 3D growth contributed to the wild type pool.

### Isolation of genomic DNA and sequencing

Genomic DNA was extracted from 7 d old protonemata grown on cellophane overlaid BCDAT medium using the CTAB method [41]. Tissues were first ground in liquid nitrogen, suspended in 500 µl CTAB buffer (1.5% CTAB, 1.05 M NaCl, 75 mM Tris-HCl, 15 mM EDTA pH8.0) and incubated at 65°C for 10 min. An equal volume of chloroform:isoamylalcohol (24:1) was added to the suspension, which was vortexed and centrifuged at 13,000 rpm for 10 min. The upper aqueous layer was transferred to a fresh tube, mixed with 0.7 volumes of isopropanol and centrifuged at 13,000 rpm for 10 min. Pellets were then washed with 70% ethanol, allowed to air dry and resuspended in 30-50 µl sterile water. Genomic DNA was extracted from individuals and then pooled in equimolar amounts to generate the mutant (120 individuals, 3 µg total) and wild type pools (120 individuals, 3 µg total). DNA samples were sequenced using a BGISEQ-500 platform (100 bp PE read lengths) at the Beijing Genomics Institute (BGI).

### Read processing and variant calling

Raw reads were obtained from BGI and subjected to quality filtering using Trimmomatic [42] to eliminate contaminants, in addition to low quality bases and read pairs. Common Illumina adaptors were identified within sequences using Trimmomatic and the read processing settings LEADING:20 TAILING:20 SLIDINGWINDOW:5:20 MINLEN:50. BWA-MEM [43] was used to map the trimmed quality filtered paired-end read libraries to the *P. patens* genome (version Ppatens_318_v3) obtained from Phytozome V12. Any duplicated reads were removed, mapped reads were realigned around insertions or deletions, and variants called according to best practice guidelines from GATK using GATK v3.6 [44].

### Candidate gene discovery through bulk segregant analysis

A previous comparison of the sequence data from the Gd::GFP and Vx::mCherry parental strains identified a set of SNPs that distinguished the two strains [20, 23]. Bulk segregant analysis was carried out by mapping read from both wild type and mutant pools to the *P. patens* reference genome. The allele frequencies were calculated by performing comparative analyses of the strain-distinguishing variants in both pools. Strain-distinguishing variants were used for analysis if: 1) the coverage depth in both the wild type and mutant pools was > 0.5 and < 2 times the mean coverage depth and: 2) the variant quality was > 500. Allele frequencies were used to identify the chromosomal region linked to the mutant allele. To confirm the presence of the premature termination codon in the PpNOG2 gene in the Ppnog2-R mutant, the full-length transcript was amplified using the primers NOG2StSalI and NOG2-StopHindIII (**Data S2**), sequenced and compared to the wild type version.

### *Physcomitrium patens* transformation

Prior to transformation, 10-15 µg of plasmid DNA (pAct:*PpNOG2*-mGFP) was linearised using KpnI and purified. All of the solutions required for the transformation procedure were prepared immediately before commencing transformation. Initially, 2g polyethylene glycol 6000 (previously autoclaved in a flat-bottom autoclavable vial) was melted in a microwave. 5 ml filter sterilised mannitol/Ca(NO_3_)_2_ solution (7.2% mannitol, 100 mM Ca(NO_3_)_2_, 10 mM Tris pH8.0) was then added to the molten PEG and allowed to incubate at room temperature until required (approx. 2 h). Protoplasts were isolated from *nog2-R* (or Gransden wild type) as described above in *‘UV mutagenesis and screening’*. Following cell density determination using a haemocytometer, protoplasts were resuspended in MMM (0.5 M mannitol, 15 mM MgCl_2_, 0.1% MES pH5.6) to achieve a final cell density of 1.5×10^6^ cells mL^-1^. 10-15 µg of linearised construct was added to a round bottomed tube (non-linearised DNA was used for transient transformations). 300 µL protoplast suspension and 300 µL PEG solution were then added to the tubes in drops and mixed by gently tilting the tubes. The samples were heat-shocked in a waterbath for 5 min at 45°C and incubated for an additional 5 min at room temperature. 300 µL 8% mannitol was added to each tube five times at 4-6 min intervals with mixing after each addition. 1 mL 8% mannitol was then added to each tube five times at 4-6 min intervals with mixing after each addition. The tubes were then centrifuged for 3 min at 120 xg at room temperature with no braking to pellet the protoplasts. For transient transformations, protoplasts were resuspended in liquid BCD medium containing 10 mM CaCl_2_, 8% mannitol and 0.5% glucose. Tubes were then wrapped in aluminium foil to protect them from light and incubated overnight at 24°C. For stable transformations, protoplasts were resuspended gently in 3 mL 8% mannitol. 1 mL of the protoplast suspension was then added to each of three cellophane overlaid plates containing BCDATG medium. Protoplasts were incubated at 24°C in the dark for 24-48 h, and then grown under standard conditions for one week. Cellophane discs were then transferred to BCDAT plates supplemented with antibiotics and grown for one week. Cellophane discs were then transferred to BCDAT (without antibiotics) and grown for a further week. Using forceps, regenerating lines were then transferred to BCD medium supplemented with antibiotics for longer term growth.

### Generation of *Ppnog2-R* complementation lines

The full-length *PpNOG2* coding sequence (excluding the stop codon) was amplified from *P. patens* Villersexel wild-type protonemata cDNA using the primers NOG2StSalI and NOG2- StopHindIII and ligated into SalI/HindIII cut pZAG1 plasmid (a gift from Yuji Hiwatashi). The resultant construct, pAct:*PpNOG2*-mGFP (**Figure S4**), was linearised using KpnI and transformed into protoplasts isolated from *Ppnog2-R* mutants. Stable transformants were selected using 100 µgmL^-1^ Zeocin. Primer sequences are listed in (**Data S2**).

### Quantitative RT-PCR

Total RNA was isolated using the RNeasy Mini kit (QIAGEN). 1 µg RNA was DNase treated (Turbo DNase, Ambion) and cDNA was subsequently synthesised using Superscript III Reverse Transcriptase (Invitrogen). Primer pairs were designed to amplify ∼100-150bp fragment of each of the genes of interest. Genes examined included *PpDEK1, PpSHR1, PpSHR2, PpSCR1, PpSCR2, PpSCR3, PpIMK3A, PpCLE1, PpCLE2, PpCLE3, PpCLE4, PpCLE5, PpCLE6, PpCLE7, PpCLV1A, PpCLV1B* and *PpRPK2* (for primer sequences see **Data S2**). Transcripts for the following genes were undetectable in wild type, and in both *Ppnog1-R* and *Ppnog2-R* mutants and were eliminated from further studies: *PpSHR1, PpSHR2, PpSCR3, PpCLE1, PpCLE2, PpCLE6, PpCLV1A* and *PpCLV1B*. Amplification was detected on a StepOne™ Real-Time PCR System (Applied Biosystems) using SYBR Green (Life Technologies). Cycling conditions comprised 95°C for 5 min, and 40 cycles of 95°C for 15 s and 60°C for 1 min. Minus RT controls were checked for genomic DNA contamination using control primer pairs. To test whether *PpNOG2* expression was affected by either auxin and/or cytokinin treatment, 7 d old wild type Vx protonemata (grown on cellophane overlaid BCD) was transferred to BCD medium supplemented with 100 nm BAP, 100 nm NAA or 100 nm BAP/100 nm NAA. Controls (no BAP or NAA) were also included. Tissues were grown for 3 d and then material was harvested for RNA isolation. For all other experiments, RNA was isolated from protonemata grown on cellophane overlaid BCD medium under standard conditions for 12 d.

### Phylogenetic analysis

The complete set of predicted proteomes for Arabidopsis thaliana, *Oryza sativa, Selaginella moellendorffii, Marchantia polymorpha, Physcomitrium patens* (formerly *Physcomitrella patens*), *Ostreococcus lucimarinus*, and *Micromonas pusilla* in Phytozome version 12.1 were subject to orthogroup inference using Orthofinder [45, 46]. The orthogroup containing the *PpNOG2* gene was identified and the constituent sequences aligned using MEGA X [47]. Several of the gene models for *Selaginela moellendorffii* and *Marchantia polymorpha* were checked, manually corrected and added to the alignment. The alignment was subjected to initial testing using IQTREE [48]. Failed sequences were then removed from the alignment. The best-fitting model parameters (Dayhoff) were estimated from the edited alignment and a consensus phylogenetic tree was estimated from 1000 bootstrap replicates. The data were imported into ITOL [49] to generate the pictorial representation. The phylogenetic tree for the SCR genes was constructed in the same manner.

## QUANTIFICATION AND STATISTICAL ANALYSES

### Experimental design, sampling and statistical methods

qRT-PCR experiments were carried out using three technical replicates for each of three independent biological samples, alongside water controls. Ct values were calculated from raw amplification data using the Real-time PCR Miner software (http://www.ewindup.info/miner/). The mean Ct value between the three technical replicates was then calculated. Fold changes in gene expression were calculated relative to controls using the 2-ΔCT method [50]. Two genes were used as constitutive controls: EF1alpha (Pp1s84_186V6) and ACT7 (Pp1s198_154V6). An ANOVA test (or t-test if appropriate) was performed to determine whether relative expression changes were statistically significant. A phenotypic segregation analysis was performed on the progeny derived from a single *Ppnog2-R*/Gd hybrid line. 660 individuals were phenotypically scored, and a Chi-square test was performed to check for alignment with the expected 1:4:1 segregation ratio.

## DATA AND SOFTWARE AVAILABILITY

The accession number for the raw genome sequence reads for the Gd::GFP and Vx::mCherry parental lines is EBI ArrayExpress: E-MTAB-5096. The accession number for the raw genome sequence reads reported in this paper is EBI ArrayExpress: E-MTAB-7502. The accession numbers for the sequences of full length *PpNOG2* cDNA plus the two different splice variants in *nog2-R* mutants reported in this paper are NCBI: MT266984, MT266985, and MT266986, respectively.

## SUPPLEMENTAL INFORMATION

**Figure S1. Bulk segregant analysis via somatic hybridization** (A) *Ppnog2-R* and Gd::GFP haploid lines and a *Ppnog2-R*/Gd::GFP diploid line. Images show green fluorescent protein (GFP, green), mCherry (red), chlorophyll autofluorescence (blue) and a merge of all four. Scale = 25 µm. (B, C) Allele frequency plots for *Ppnog2-R* (B) and WT (C) segregants on chromosome 2 of *P. patens*.

**Figure S2. Generation of *PpNOG2* complementation lines** Schematic diagram of the pAct::PpNOG2-GFP construct that was used to complement

*Ppnog2-R* mutants. The construct was targeted to the 213 locus as shown.

**Figure S3. The PpNOG2 protein localizes at or near to the plasma membrane**

A) Transient expression of pAct::PpNOG2-GFP in P. patens protoplasts. Images show green fluorescent protein (GFP, green), chlorophyll autofluorescence and a merge of both. Scale = 10 µm. B) Schematic of the PpNOG2 protein topology as predicted by TMpred (see **Data S2** for further details). Three transmembrane (TM) domains are predicted to exist, in addition to an extracellular loop region. The putative location of conserved aspartic acid (D) and histidine (H) have been indicated.

**Figure S4. Maximum-likelihood phylogenetic analysis of PpNOG2-related proteins**

The tree was rooted using chlorophyte sequences, and numbers correspond to bootstrap values. PpNOG2 (Pp3c2_29140) has been highlighted in red.

**Figure S5. Previous model for 3D growth regulation in *P. patens***

Previously proposed model for 3D growth regulation suggests that PpNOG1 and PpDEK1 operate antagonistically to regulate the transcription of PpAPB genes, and then cooperate with CLAVATA to deliver correct cytokinin signalling outputs; prevent ectopic initiation of buds and ensure division planes are correctly oriented.

**Figure S6. Maximum-likelihood phylogenetic analysis of SCR-related proteins**

The tree was rooted to *P. patens* sequences, which form an outgroup. Bootstrap values are shown on branches. Green shading indicates the AtSCL23 clade and yellow shading indicates the AtSCR clade.

## Notes

### Competing Interest Statement

The authors have declared no competing interest.

