## Supplementary figures and images for "*NO GAMETOPHORES 2* is a novel regulator of the 2D to 3D growth transition in the moss *Physcomitrium patens*"

### Figure S1

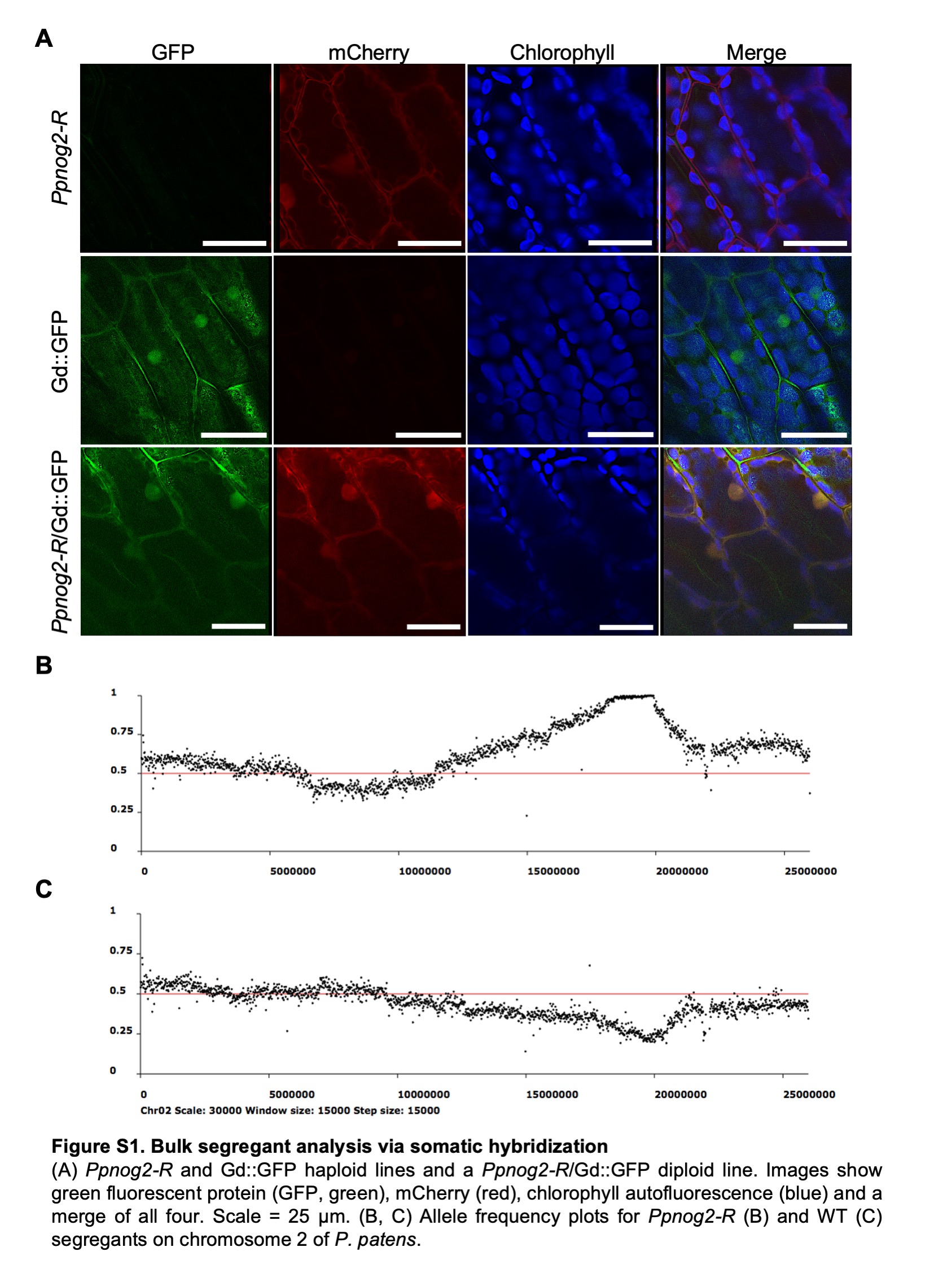

### Figure S2

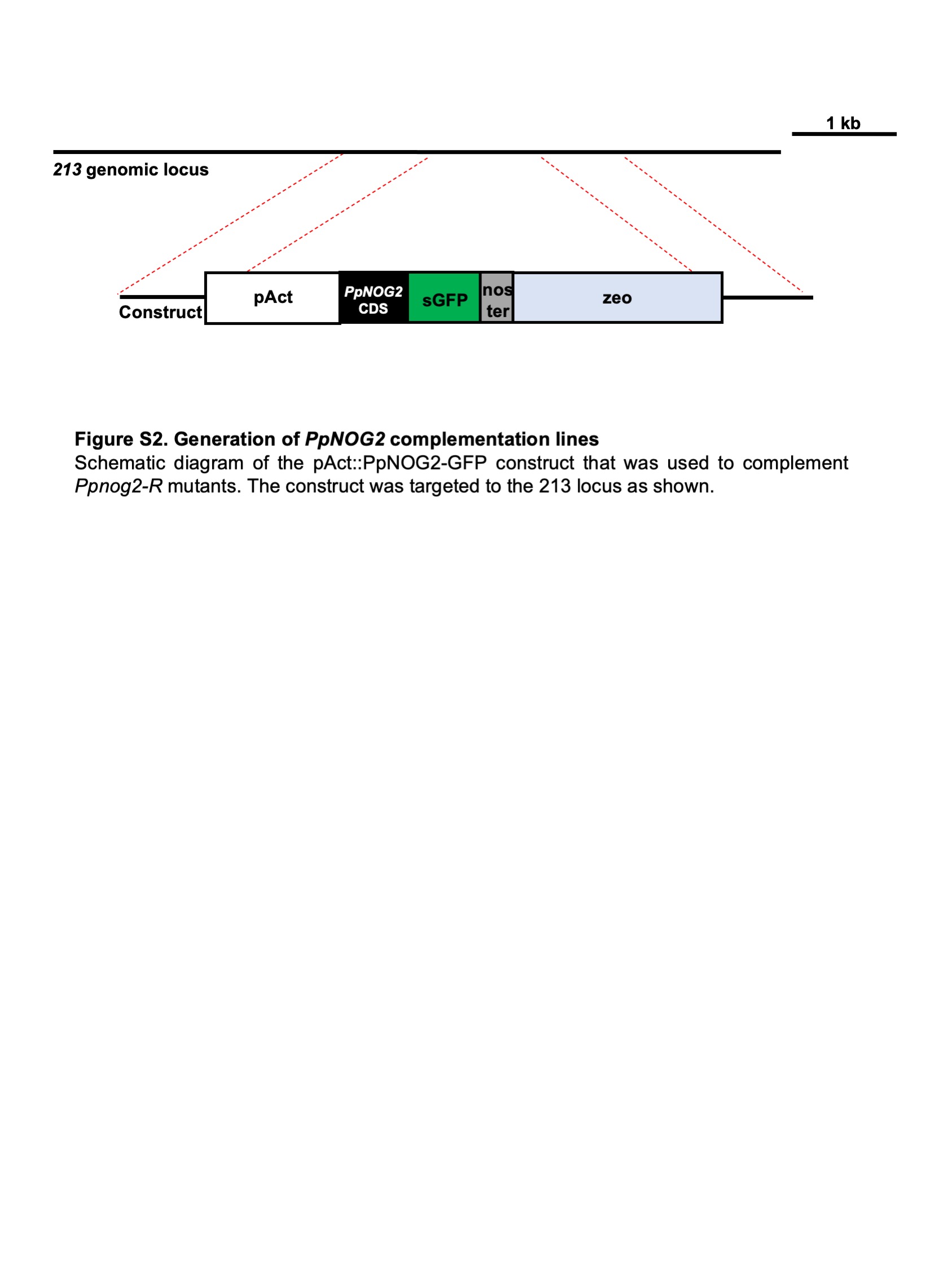

### Figure S3

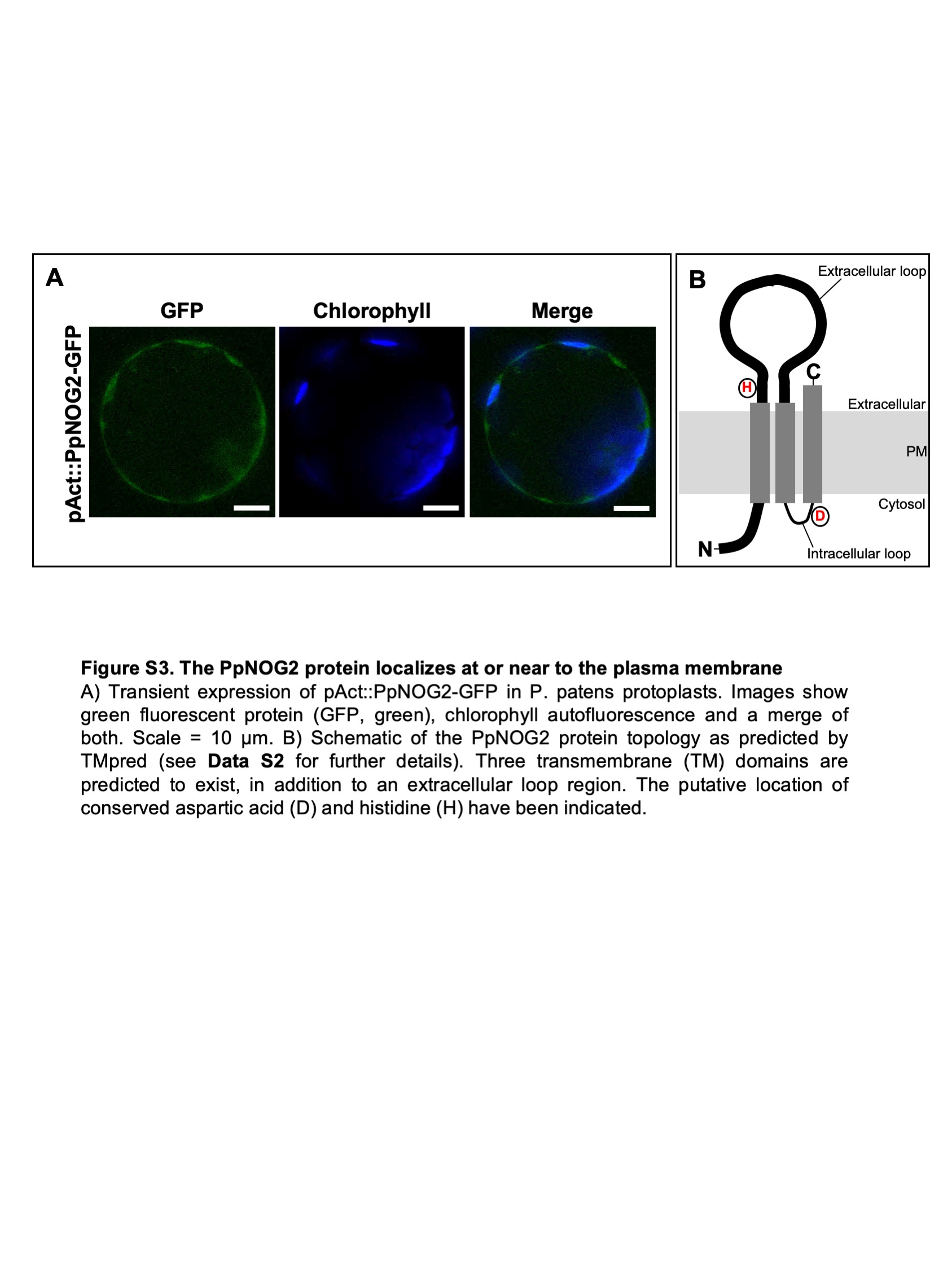

### Figure S4

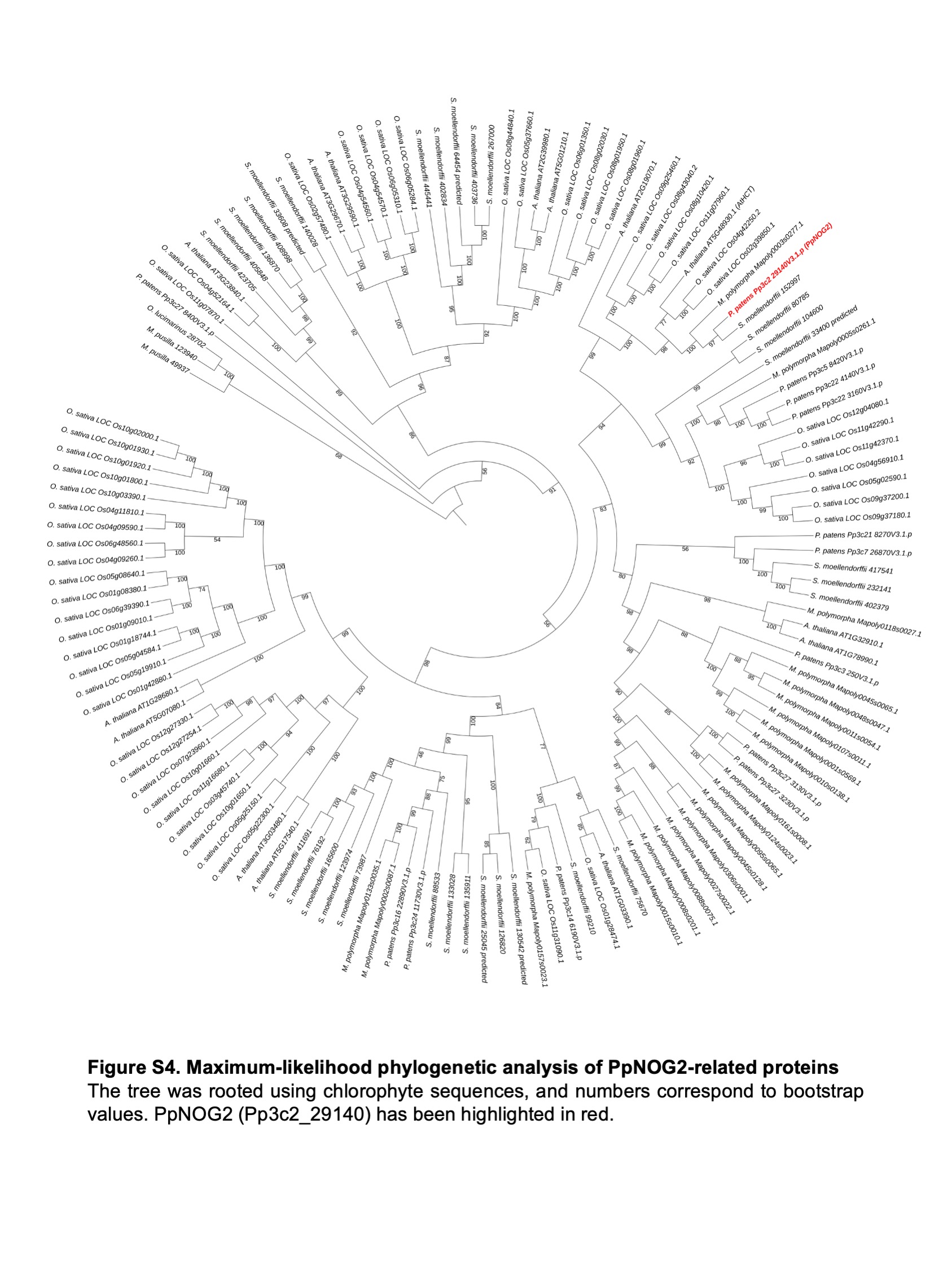

### Figure S5

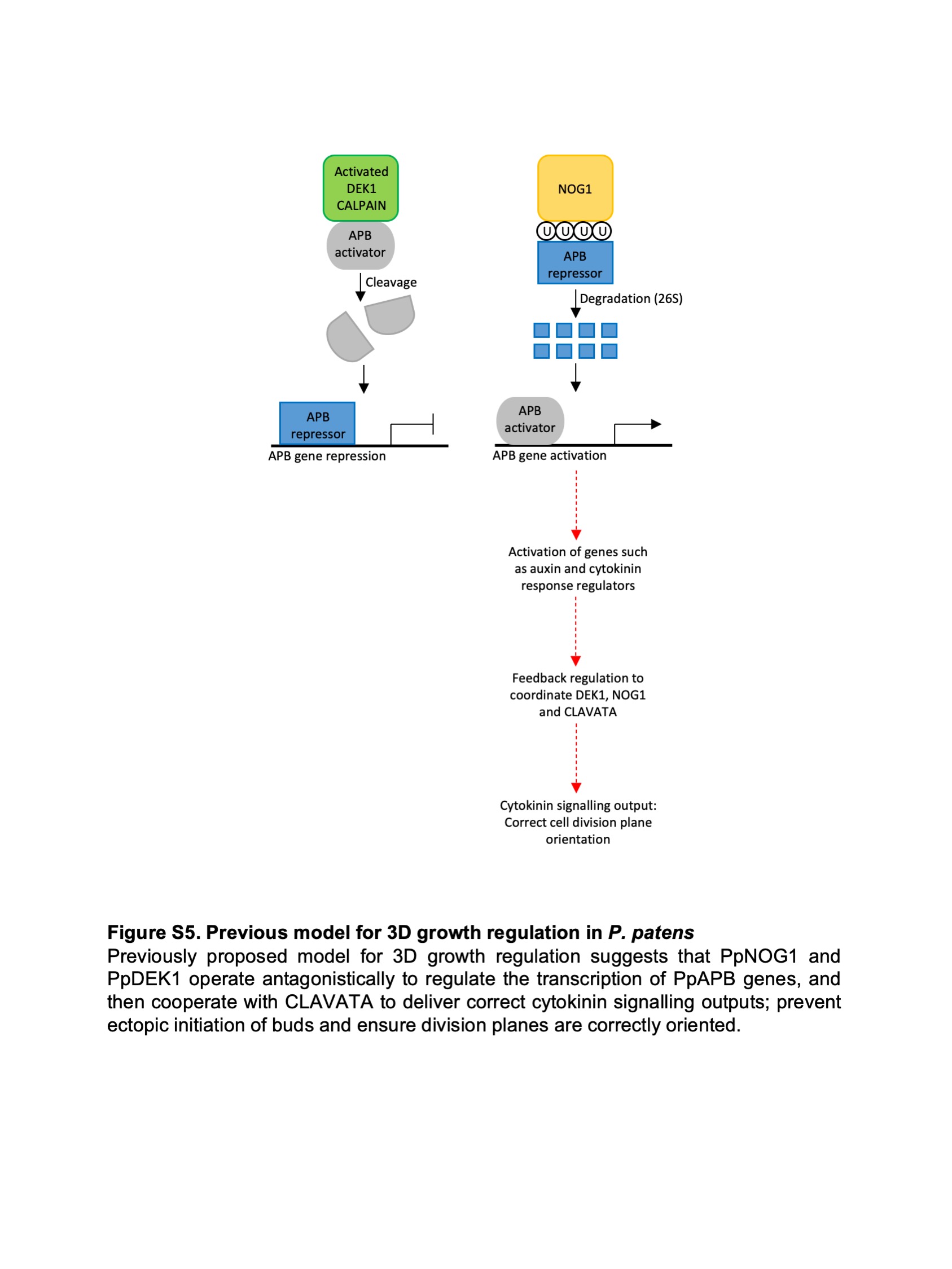

### Figure S6

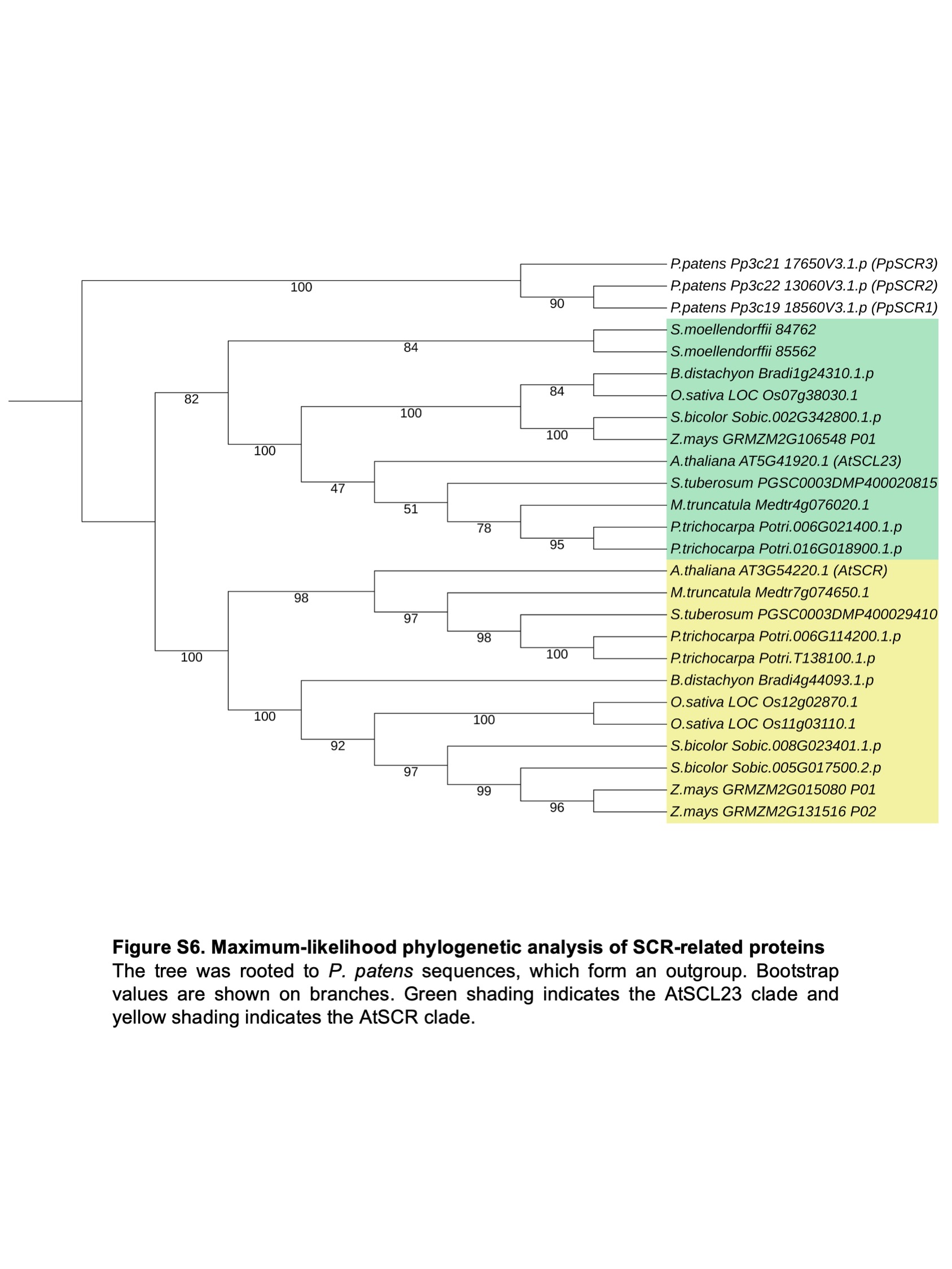
