## Supplementary material for "*NO GAMETOPHORES 2* is a novel regulator of the 2D to 3D growth transition in the moss *Physcomitrium patens*": Data S1

```
Pp3c2_29140V3.1      MAAASQVQLDVTIRKQSMVRPSEPTPERILWNSGLDLVIPRIHTQSVYFYNNNDGSDDFL	60
AT5G48930.1          --------MKINIRDSTMVRPATETPITNLWNSNVDLVIPRFHTPSVYFYRPT-GASNFF	51
                             :.:.**..:****:  **   ****.:******:** *****. . *:.:*:

Pp3c2_29140V3.1      NHEKLAEALGKTLVPFYPMAGRLKRGEGGRVEINCNGEGVLLVEAEADAKITDFGEFAPD	120
AT5G48930.1          DPQVMKEALSKALVPFYPMAGRLKRDDDGRIEIDCNGAGVLFVVADTPSVIDDFGDFAPT	111
                     : : : ***.*:*************.:.**:**:*** ***:* *:: : * ***:***

Pp3c2_29140V3.1      PRFRHLVPQVDYSQEISSFPLLVLQITFFKCGGASLGVGMQHHVADGMSGLHCVNTWSEM	180
AT5G48930.1          LNLRQLIPEVDHSAGIHSFPLLVLQVTFFKCGGASLGVGMQHHAADGFSGLHFINTWSDM	171
                      .:*:*:*:**:*  * ********:*****************.***:**** :****:*

Pp3c2_29140V3.1      ARGIPLKVKPFIDRTLLKANSPPRPMFPHIEYQPPPRLMKPENSKSNGNGHSNGHSNGLA	240
AT5G48930.1          ARGLDLTIPPFIDRTLLRARDPPQPAFHHVEYQPAPSMKIPLDPSKSGPE----------	221
                     ***: *.: ********:*..**:* * *:**** * :  * : ...*

Pp3c2_29140V3.1      NGHANVHANGLANGHANGHSKGHANNYSNGHHTNGMEITNGAFHTNGNGNGVVNDSHVAN	300
AT5G48930.1          ------------------------------------------------------------	221


Pp3c2_29140V3.1      RKSNGAHMENGNGNGNGDFNENHEKQGSGNGNSVCAGGGHNHANGNENQSAVQLNVKKMT	360
AT5G48930.1          ------------------------------------------------------------	221


Pp3c2_29140V3.1      NGVHNSNGSTNEEAKDEDLPMAVRVFRFTKEQLATLKRMAVEEKADVTFSSYEMLSGHIW	420
AT5G48930.1          -------------------NTTVSIFKLTRDQLVALKAKSKEDGNTVSYSSYEMLAGHVW	262
                                          :* :*::*::**.:**  : *:   *::******:**:*

Pp3c2_29140V3.1      KCITQARKLAESQETKLFVATDGRSRLNPPLPKGYFGNVIFTCTPIATAGELVSNPITYA	480
AT5G48930.1          RSVGKARGLPNDQETKLYIATDGRSRLRPQLPPGYFGNVIFTATPLAVAGDLLSKPTWYA	322
                     :.: :** * :.*****::********.* ** *********.**:*.**:*:*:*  **

Pp3c2_29140V3.1      ARKIHDSLARMNDEYLRSALDYLETQEDISKLVRGAHHFNSPNLGITSWARMPTYDCDFG	540
AT5G48930.1          AGQIHDFLVRMDDNYLRSALDYLEMQPDLSALVRGAHTYKCPNLGITSWVRLPIYDADFG	382
                     * :*** *.**:*:********** * *:* ****** ::.********.*:* **.***

Pp3c2_29140V3.1      WGRPIFMGPATIAYEGLVYVLASPVNDGSLSLSLGLRSDHMDTFAKLVASF*	591
AT5G48930.1          WGRPIFMGPGGIPYEGLSFVLPSPTNDGSLSVAIALQSEHMKLFEKFLFEI*	433
                     *********. * **** :** **.******:::.*:*:**. * *:: .:*

Active site (Levsh et al., 2016)
Predicted TM domain (TMpred)
```
