## Supplementary material for "*NO GAMETOPHORES 2* is a novel regulator of the 2D to 3D growth transition in the moss *Physcomitrium patens*": Data S2

**Cloning and sequencing primers:**

**Generation of *Ppnog2-R* complementation lines**

| | | |
|---|---|---|
| NOG2StSalI | aaagtcgacATGGCCGCCGCAAGTCAAGTT | Amplification (and sequencing) of full-length *PpNOG2* transcript - Forward primer |
| NOG2-StopHindIII | aaaaagcttGAAGGATGCCACTAGTTTGG | Amplification (and sequencing) of full-length *PpNOG2* transcript - Reverse primer |

**qPCR primers:**

| | | |
|---|---|---|
| PpACT7qPCRF2 | GGGGAAGGTGAGGATGTCC | Real-time PCR amplification of *PpACT7* (Pp3c3_33410V3.1) - Forward primer |
| PpACT7qPCRR2 | GATGCTCGGAAACACGGC | Real-time PCR amplification of *PpACT7* (Pp3c3_33410V3.1) - Reverse primer |
| EF1alphaqPCRF | AATCATACATTTCACCTCGCC | Real-time PCR amplification of *PpEF1A* (Pp3c2_10310V3.1) - Forward primer |
| EF1alphaqPCRR | GATCAGTGGGTAGAAGTGAC | Real-time PCR amplification of *PpEF1A* (Pp3c2_10310V3.1) - Reverse primer |
| PpNOG1qPCR_F2 | TTTAGAGCGTCGCATCCTTG | Real-time PCR amplification of *PpNOG1* (Pp3c1_11420V3.1) - Forward primer |
| PpNOG1qPCR_R2 | CGTCGCATCTTCCACTTCAG | Real-time PCR amplification of *PpNOG1* (Pp3c1_11420V3.1) - Reverse primer |
| NOG2_29140_qPCRF1 | CACCTCGTTCCTCAAGTGGA | Real-time PCR amplification of *PpNOG2* (Pp3c2_29140V3.1) - Forward primer |
| NOG2_29140_qPCRR1 | CATGCCCACTCCTAGCGA | Real-time PCR amplification of *PpNOG2* (Pp3c2_29140V3.1) - Reverse primer |
| PpAPB1GSP-F1 | CCATCCACGCGGTTGATAGT | Real-time PCR amplification of *PpAPB1* (Pp3c15_24980V3.1) - Forward primer |
| PpAPB1GSP-R1 | TCACAGGATCACGAAGGACAAA | Real-time PCR amplification of *PpAPB1* (Pp3c15_24980V3.1) - Reverse primer |
| PpAPB2GSP-F1 | CGGTCCGCGGGAAAG | Real-time PCR amplification of *PpAPB2* (Pp3c9_25570V3.1) - Forward primer |
| PpAPB2GSP-R1 | TGGGACTGGGAACTCGTCAT | Real-time PCR amplification of *PpAPB2* (Pp3c9_25570V3.1) - Reverse primer |
| PpAPB3GSP-F1 | GGCGAATTGTCGGCATCT | Real-time PCR amplification of *PpAPB3* (Pp3c9_25400V3.1) - Forward primer |
| PpAPB3GSP-R1 | TCTGCGCCTGACCTGAGTACT | Real-time PCR amplification of *PpAPB3* (Pp3c9_25400V3.1) - Reverse primer |
| PpAPB4qPCR_F3 | TTGCAGTTGTAACCTGACGG | Real-time PCR amplification of *PpAPB4* (Pp3c15_24790V3.1) - Forward primer |
| PpAPB4qPCR_R3 | AAGGGCCAAAGGAGGGAATT | Real-time PCR amplification of *PpAPB4* (Pp3c15_24790V3.1) - Reverse primer |
| PpDEK1a_qPCRF1 | GCTTCTGTTGGGTGCAATCT | Real-time PCR amplification of *PpDEK1* (Pp3c17_17550V3.1) - Forward primer |
| PpDEK1a_qPCRR1 | GTTATGCGACGCCCAATGAT | Real-time PCR amplification of *PpDEK1* (Pp3c17_17550V3.1) - Reverse primer |
| PpSHR1_qPCRF1 | CGATGCCACCGAAGTCTAAC | Real-time PCR amplification of *PpSHR1* (Pp3c3_4950V3.1) - Forward primer |
| PpSHR1_qPCRR1 | CAAAAGTTGTCCAGGGGCTC | Real-time PCR amplification of *PpSHR1* (Pp3c3_4950V3.1) - Reverse primer |
| PpSHR2_qPCRF1 | GCGGAGAGGACCTATTCGTT | Real-time PCR amplification of *PpSHR2* (Pp3c7_24880V3.1) - Forward primer |
| PpSHR2_qPCRR1 | AAGGCCTCCATCATAGCTCC | Real-time PCR amplification of *PpSHR2* (Pp3c7_24880V3.1) - Reverse primer |
| PpSCR1_qPCRF1 | TCGTGACTAAGGAACCCACC | Real-time PCR amplification of *PpSCR1* (Pp3c19_18560V3.1) - Forward primer |
| PpSCR1_qPCRR1 | CCGCTGTGAAAAGGATCTCG | Real-time PCR amplification of *PpSCR1* (Pp3c19_18560V3.1) - Reverse primer |
| PpSCR2_qPCRF1 | TTGAGCAAGATTTCCGCCAC | Real-time PCR amplification of *PpSCR2* (Pp3c22_13060V3.1) - Forward primer |
| PpSCR2_qPCRR1 | GTTGCTGCTCGACCATGTAG | Real-time PCR amplification of *PpSCR2* (Pp3c22_13060V3.1) - Reverse primer |
| PpSCR3_qPCRF1 | TGTTCATAGAAGAGCAGATTCGC | Real-time PCR amplification of *PpSCR3* (Pp3c21_17650V3.1) - Forward primer |
| PpSCR3_qPCRR1 | CGAGGCCACTGGAGAAGAA | Real-time PCR amplification of *PpSCR3* (Pp3c21_17650V3.1) - Reverse primer |
| PpIMK3A_qPCRF1 | CCCCTGAGCTGACCAAACT | Real-time PCR amplification of *PpIMK3A* (Pp3c3_28520V3.1) - Forward primer |
| PpIMK3A_qPCRR1 | TCAATGGCTCCGTCAGTAGT | Real-time PCR amplification of *PpIMK3A* (Pp3c3_28520V3.1) - Reverse primer |
| PpCLE1_qPCRF1 | GGTCAGCTGGAGTTTTGCAC | Real-time PCR amplification of *PpCLE1* (Pp3c7_11040V3.1) - Forward primer |
| PpCLE1_qPCRR1 | TGCGTTCGTATGTCTCTCTGA | Real-time PCR amplification of *PpCLE1* (Pp3c7_11040V3.1) - Reverse primer |
| PpCLE2_qPCRF1 | TGCTGCTGGTACTCACGTTA | Real-time PCR amplification of *PpCLE2* (Pp3c1_13720V3.1) - Forward primer |
| PpCLE2_qPCRR1 | TTCTCAACCGCATCTGAACG | Real-time PCR amplification of *PpCLE2* (Pp3c1_13720V3.1) - Reverse primer |
| PpCLE3_qPCRF1 | GCTCACTCTGGCATCACTTG | Real-time PCR amplification of *PpCLE3* (Pp3c3_10020V3.1) - Forward primer |
| PpCLE3_qPCRR1 | TCGATCACAGCACTTGAGGT | Real-time PCR amplification of *PpCLE3* (Pp3c3_10020V3.1) - Reverse primer |

| | | |
|---|---|---|
| PpCLE4_qPCRF1 | TCAACAACTCGCGACACTTG | Real-time PCR amplification of *PpCLE4* (Pp3c26_11430V3.1) - Forward primer |
| PpCLE4_qPCRR1 | GACAAGCCTACGAAACCGAC | Real-time PCR amplification of *PpCLE4* (Pp3c26_11430V3.1) - Reverse primer |
| PpCLE5_qPCRF1 | TGGAGGACGATGGGAGAAAC | Real-time PCR amplification of *PpCLE5* (Pp3c22_4590V3.1) - Forward primer |
| PpCLE5_qPCRR1 | TCTGCCTCCACATCCCAAAT | Real-time PCR amplification of *PpCLE5* (Pp3c22_4590V3.1) - Reverse primer |
| PpCLE6_qPCRF1 | GAATCCCTCTACTCCGCGAA | Real-time PCR amplification of *PpCLE6* (Pp3c19_6950V3.1) - Forward primer |
| PpCLE6_qPCRR1 | CAAATCGGGTCCACAGCATC | Real-time PCR amplification of *PpCLE6* (Pp3c19_6950V3.1) - Reverse primer |
| PpCLE7_qPCRF1 | GATTCTGAGCTTGAGTGCCG | Real-time PCR amplification of *PpCLE*7 (Pp3c21_5600V3.1) - Forward primer |
| PpCLE7_qPCRR1 | TTCTGGCTCGACAAAGACCT | Real-time PCR amplification of *PpCLE*7 (Pp3c21_5600V3.1) - Reverse primer |
| PpCLV1A_qPCRF1 | CCTCATGCTGCAATACGTGA | Real-time PCR amplification of *PpCLV1A* (Pp3c13_13360V3.1) - Forward primer |
| PpCLV1A_qPCRR1 | TCGTTGTTGAAGCAGTCCAG | Real-time PCR amplification of *PpCLV1A* (Pp3c13_13360V3.1) - Reverse primer |
| PpCLV1B_qPCRF1 | CGGCTTGAACGGAAACTCTC | Real-time PCR amplification of *PpCLV1B* (Pp3c6_21940V3.1) - Forward primer |
| PpCLV1B_qPCRR1 | TGGGATGCTCGAACTGAAGT | Real-time PCR amplification of *PpCLV1B* (Pp3c6_21940V3.1) - Reverse primer |
| PpRPK2_qPCRF1 | GATCGACCGTCCTCTCCAAT | Real-time PCR amplification of *PpRPK2* (Pp3c7_5570V3.1) - Forward primer |
| PpRPK2_qPCRR1 | GCCAAGACGCCGTACAATAG | Real-time PCR amplification of *PpRPK2* (Pp3c7_5570V3.1) - Reverse primer |
